# Gene copy number variation (gCNV) contributes to adaptation along environmental gradient

**DOI:** 10.1101/2025.09.12.675866

**Authors:** Qiujie Zhou, Martin Lascoux, Pascal Milesi

## Abstract

Gene copy number variations (gCNVs) are structural variations that represent a significant source of genetic polymorphism. While single nucleotide polymorphisms (SNPs) have been the primary focus of population and quantitative genomics, recent studies indicated that gCNVs could also play an important role in adaptation notably because of their multiallelic and quantitative nature. In this study, we investigate the role of gCNVs in local adaptation along environmental gradients using extensive genomic datasets in Norway spruce (*Picea abies*) and Siberian spruce (*P. obovata*). We used a robust pipeline for the detection and quantification of gCNVs from short-read exome capture data and used haploid samples for validation. We showed that gCNVs are pervasive, representing approximately 11% of the protein coding genes and are notably enriched in genes involved in response to environmental stress, such as temperature tolerance, immune response, and metal ion regulation. Population genetic structure at gCNV was similar to that observed at SNPs. However, some gCNVs also display distinctive adaptive signatures not captured by SNPs. Finally, we conducted gCNV-based genotype-environment association (GEA) and genome-wide association studies (GWAS) to further evidence that gCNVs contribute to local adaptation patterns and to the control of quantitative traits.

## Introduction

Understanding the genetic basis of adaptation is a central issue in evolutionary biology and has implications for many fields, from conservation biology and population management to plant and animal breeding and health. The development of next-generation sequencing has facilitated the acquisition of population-level genomic data, providing insights into patterns of local adaptation and of its genetic control. Genome scans, genotype-environment or genotype-phenotype association studies have been widely used to characterize the genetic architecture of adaptative traits, and to relate these architectures to various factors such as the selection at the loci underlying the variation in the trait, or even the biology of the focal species (e.g., generation time, intensity of gene flow, mating system).

In organisms occupying large geographical ranges, such as forest trees, local adaptation often occurs along environmental gradients. The source of selection can thus be highly dimensional, encompassing abiotic and biotic factors (e.g. climatic variables, photoperiod, soil types, presence of pathogens) and the response can affect many, often correlated, phenotypic traits. Hence, one can expect a polygenic architecture of adaptation and a quantitative relationship between environmental, genotypic and phenotypic variation (e.g. Milesi et al. 2019). To date, most studies of adaptation along environmental gradients have relied on the role of single nucleotide polymorphisms (SNPs). Yet, a main outcome of a decade of high-throughput sequencing is that genomic structural variations (SVs) encompass more genetic variability than SNPs (e.g. Catanach et al. 2019; Tigano 2020). Recent studies highlighted an important role for SVs on both long and short evolutionary timescales (Weissensteiner et al. 2020; Y. Zhou et al. 2019; Finnegan et al. 2023; Peona et al. 2022), with a large number of cases suggesting their significant contributions to local adaptation (e.g. Westram et al. 2021; Lecomte et al. 2024; Yan et al. 2021; Ben-Jemaa et al. 2024; Cayuela et al. 2021). Hence, especially in non-model species where available polymorphism often covers a limited part of the genome, it would be advantageous to include SV as additional markers.

In contrast to other types of SVs (e.g. inversions and translocations), deletions and duplications are unbalanced mutations that affect the dosage (amount) of a DNA sequence, resulting in copy number variations (CNVs). In the case of gCNV, these mutations lead to a variation in the number of copies of a gene among individuals. Gene copy number variations (gCNVs) are widespread in eukaryotic genomes (e.g. Schiessl et al. 2017; Zmienko et al. 2016; Prunier et al. 2017), supported by a gene duplication rate per gene per generation is many orders of magnitude larger than point mutation rate (e.g., Mahmoud et al. 2019; Katju and Bergthorsson 2013). While most gene duplications are probably highly deleterious in the early stages (e.g, Schrider et al. 2013), since the first likely consequence of a change in gene copy number is a change in the amount of the gene’s product (e.g. Labbé et al. 2014; Shao et al. 2019), they can also form the basis of adaptations (e.g. detoxification and resistance to xenobiotics, Kondrashov 2012 and references therein). Different gene copy numbers can thus be associated with different phenotypic value (e.g. Assogba et al. 2016; Milesi et al. 2022 in insect, Wei et al. 2023 in angiosperms) and can be selected for in different environments (e.g. Dorant et al. 2020 in crustacean, Cayuela et al. 2021 in amphibian, Kuo et al. 2024 in angiosperms). Their multiallelic and quantitative nature therefore make gCNVs natural candidates in the control of quantitative traits and adaptation along environmental gradients.

In spite of their potential as a molecular marker for evolutionary studies, genome-wide screening of gCNVs has remained confined to a limited number of organisms (e.g. Redon et al. 2006; Y. Zhou et al. 2022; Prunier et al. 2017). These genome-wide screenings have used long-reads sequencing or specially designed technologies such as array-based comparative genome hybridization (aCGH) and SNP-array (McCarroll and Altshuler 2007; Prunier et al. 2017). Furthermore, gCNVs were often subsumed under the term “structural variants” and considered as bi-allelic markers together with other structural variants such as inversions (e.g. Y. Zhou et al. 2022; Kang et al. 2023). The quantitative nature of gCNVs and their use for population and quantitative genomics studies has thus been largely overlooked (Mérot et al. 2020; Lindstedt et al. 2025; Conrad and Hurles 2007, but see Chiang et al. 2017; Sjödin and Jakobsson 2012).

Here, we test the hypothesis that gene copy number variations contribute to adaptation along environmental gradient. To do so, we investigated the contribution of gCNVs to quantitative traits and local adaptation in Norway and Siberian spruce (*Picea abies* [L.] H. Karst and *P. obovata* Ledeb., respectively). Norway and Siberian spruce are two keystone boreal forest tree species which form a syngameon (Q. Zhou et al. 2024) with a joint distribution range that extends from the Alpine Mountain range in the east to the Sea of Okhotsk in the West, spanning a large array of ecological niches (Karunaratne et al, 2024). Spruce species have giga genomes with a large fraction of repeated elements (Nystedt et al. 2013; Warren et al. 2015; Nilsson et al. 2025) and extensive gene duplications (Prunier et al. 2017; Sahli 2017). Since the seminal work of Lagercrantz and Ryman 1990 spatial variation in genetic diversity at large and small geographic scales, as well as the demographic history of Norway and Siberian spruce has been studied extensively (J. Chen et al. 2019; L. Li et al. 2022; Karunarathne et al. 2024; Q. Zhou et al. 2024; Milesi et al. 2024). Furthermore, recent studies have also identified quantitative trait loci (QTL) associated to the variation of phenotypic traits and assessed their importance for local adaptation (Milesi et al. 2019; Z. Q. Chen et al. 2021; L. Li et al. 2022; CapadorlJBarreto et al. 2021; Tiret et al. 2023; J. Chen et al. 2012; 2014).

In this study we leveraged the extensive genomic resources available for *P. abies* and *P. obovata* to investigate the role of gCNVs in local adaptation. Using an innovative approach and combining information from diploid and haploid DNA we show that gCNVs are widespread in the genomes of the two spruce species (∼12% of the 26,219 targeted genes). We then further develop population genetics metrics and quantitative genomics approaches to show that gCNVs globally follow the main population structure but exhibit distinct signatures of local adaptation that were not captured by SNPs.

## Results

### Gene CNVs can be reliably called from short-read exome capture data

Calling copy number variations is challenging, even with full genome data and good quality reference genomes. Here, we leverage diploid and haploid DNA information from an extensive exome capture dataset to detect gene copy number variations (gCNVs) in two diploid exome-capture sequencing datasets. The first diploid dataset, hereafter “*Swedish cline*” dataset (plus signs, Fig. 1 and Supplementary material 1, N = 1758) includes trees sampled along a latitudinal gradient in Sweden and the second diploid dataset, hereafter “*P. abies*-*P. obovata*” dataset (filled rounds and diamonds, Fig. 1 and Supplementary material 2, N = 542) includes individuals sampled along a large longitudinal gradient. Finally, the third dataset consist of haploid tissues (megagametophytes, diamonds, Fig. 1 and Supplementary material 3, N = 180). We implemented a series of analyses to detect gCNVs from the two diploid dataset and curate the haploid dataset for false positive detection. Our approach was conservative, systematically favoring removal of false positives at the expense of false negatives.

**Figure 1.**
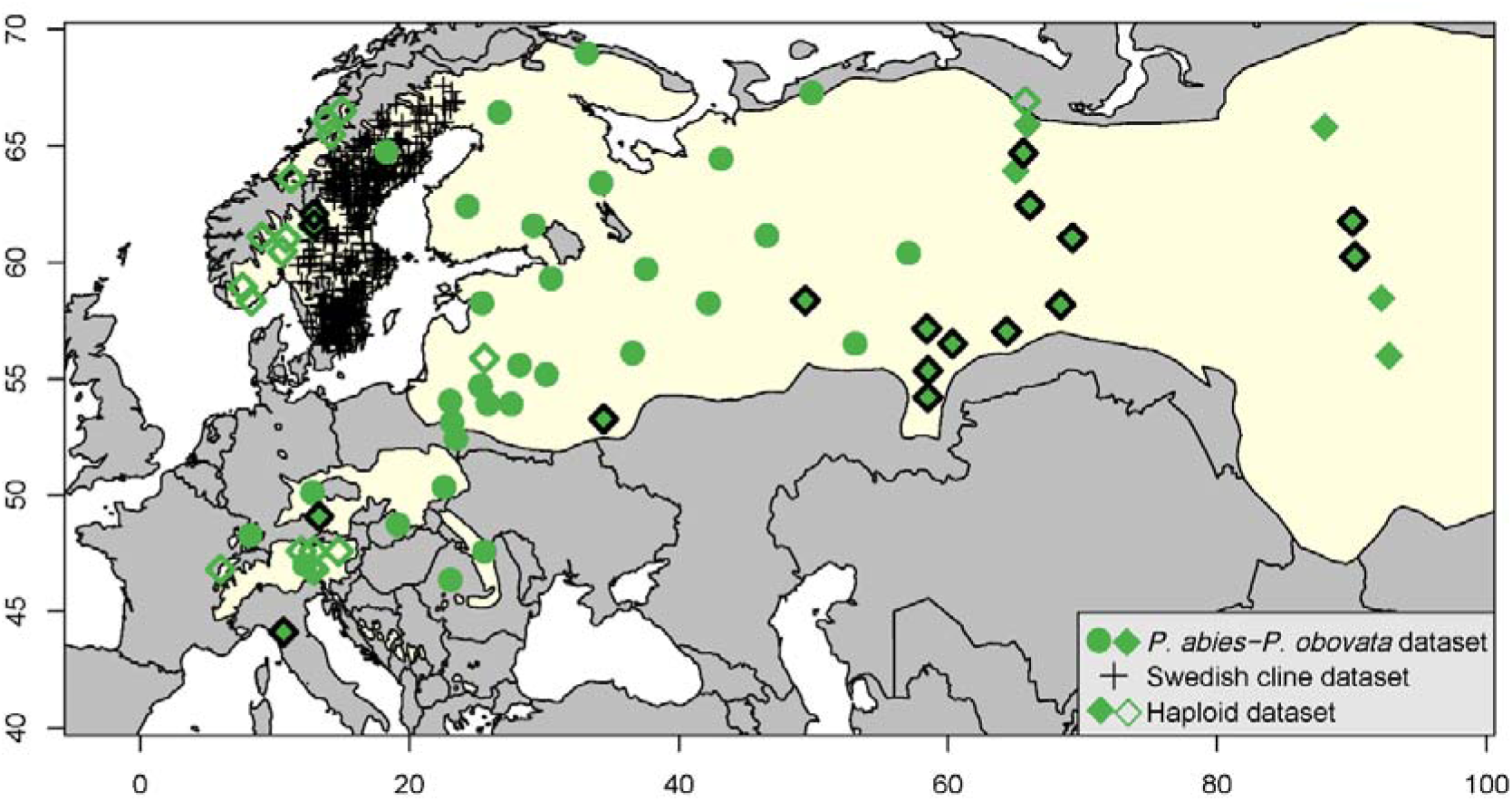
Sampling map for the three datasets included in this study. The yellow shaded area marks the joint natural distribution range of *Picea abies* and *P. obovata*. For the *P. abies-P. obovata* and the haploid dataset, location of populations sampled are indicated using filled or empty green symbols, respectively, for the *Swedish cline* dataset, individual tree locations are indicated with plus signs. Populations represented with green diamonds with thick black border indicate a subset of the *P. abies-P. obovata* dataset used for a growth chamber experiment and further genome-wide association analysis (GWAS).

First, we used the method implemented in the *rCNV* R package to identify SNPs putatively located in multi-copy regions (so called “deviant” SNPs): 46,354 (18.22% of the total number of SNPs) and 127,770 SNPs (16.90% of the total number of SNPs) were identified as such for the Swedish cline and *P. abies-P. obovata* datasets, respectively. To avoid false positives, only probes with an enrichment in deviant SNPs of at least 30% were retained as candidate markers for gene CNVs (hereafter ‘CNV-probes’). 3,460 CNV-probes (covering 3,071 genes) were identified in the Swedish cline dataset, and 3,575 CNV-probes (covering 3,041 genes) in the *P. abies-P. obovata* dataset. A total of 2,449 CNV-probes (2,173 genes) were shared by both datasets (Szymkiewicz-Simpson overlap coefficient = 0.71). After filtering out paralogs (i.e. CNV-probes with a fixed number of copies across individuals), we retained 2,801 gCNVs (encompassing 3140 probes) and 2,479 gCNVs (encompassing 2903 probes) for the Swedish cline dataset and the *P. abies-P. obovata* datasets, respectively (Szymkiewicz-Simpson overlap coefficient = 0.69). As expected for copy number variations, the genomic regions corresponding to the gCNVs showed a higher mean depth of coverage (uDoC) and a higher coefficient of variation (CV, standard deviation of DoC / square root of uDoC) than the genomic regions harboring single copy genes for both diploid datasets (Fig. 2A & B and Table 1, Welch Two Sample *t*-test, all *p*-value < 2.2e-16).

**Figure 2.**
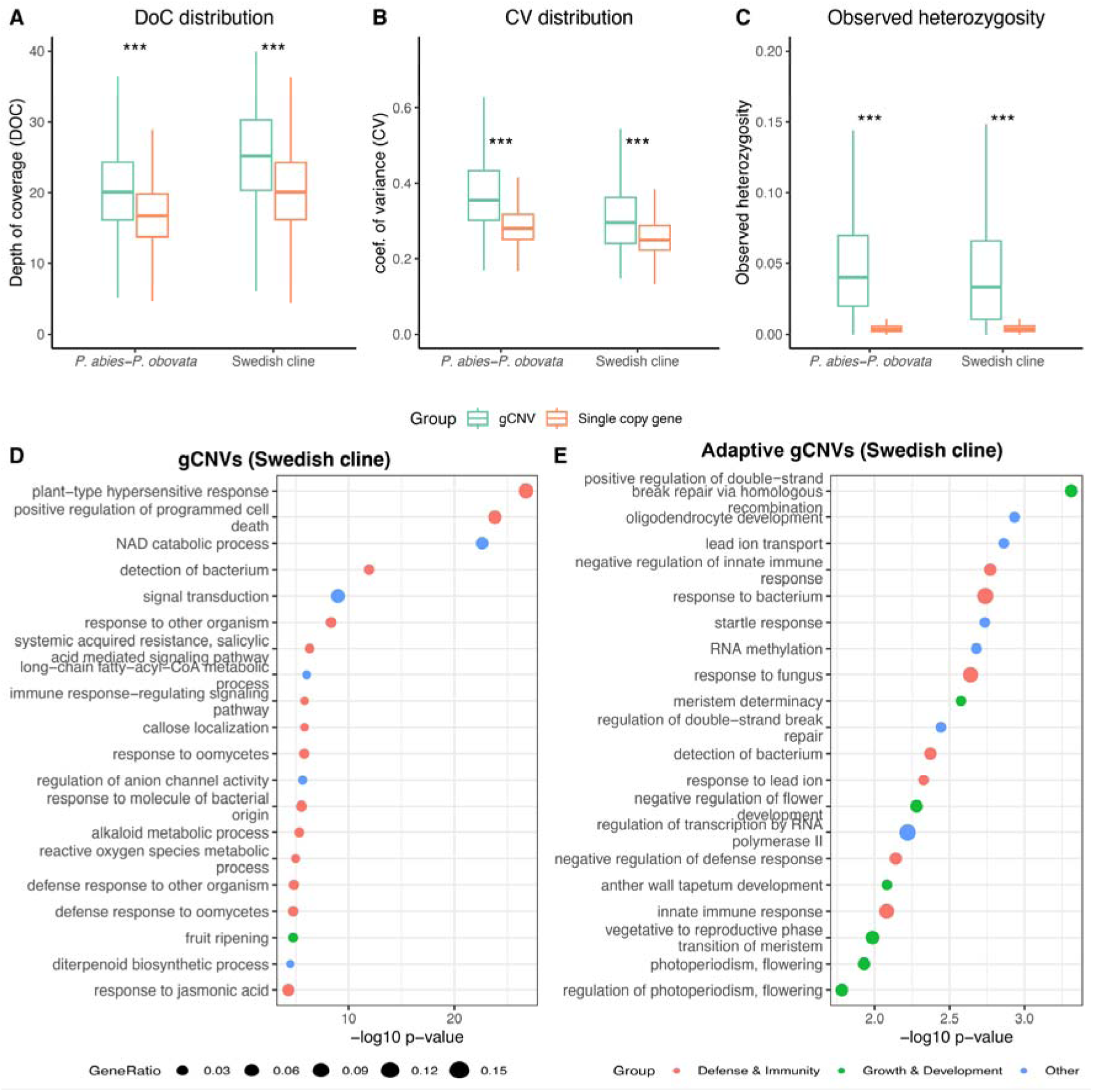
gCNVs mapping statistics and functional annotation. **A**, Depth of coverage (DoC); **B**, coefficient of variation (CV); **C**, observed heterozygosity in haploid data for genes classified as gCNVs (green) or single-copy (orange) from diploid DNA in each dataset; *** *p*-values < 2.2e^-16^. **D & E**, Gene Ontology (GO biological processes) enrichment analyses across all gCNVs (**D**) and adaptive candidate gCNVs detected through GEA for the Swedish cline datasets (**E**). Only top 20 GO terms with a *p*-value < 0.05 are shown; see Fig. S2 for the *P. abies-P.obovata* dataset, and Tables S4 and S5 for the full list of enriched GO terms.

**Table 1:** Depth of coverage statistics and observed heterozygosity for single-copy or gCNV.

|  | *Swedish Cline* |  |  |  | *P.abies - P.obovata* |  |  |  |
| --- | --- | --- | --- | --- | --- | --- | --- | --- |
|  | gCNVs | Single-copy | *t* | *df* | gCNVs | Single-copy | *t* | *df* |
| # of probes | 2801 | 21107 | - | - | 2479 | 22962 | - | - |
| $\mu\text{DoC}^{\S}$ | $28 \pm 27$ | $20 \pm 6$ | 14 | 2840 | $21 \pm 11$ | $17 \pm 5$ | 20 | 2571 |
| $\text{CV}^{\dagger}$ | $0.34 \pm 0.17$ | $0.27 \pm 0.09$ | 21 | 3027 | $0.42 \pm 0.25$ | $0.30 \pm 0.10$ | 23 | 2558 |
| $\mu\text{Ho}^{\ddagger}$ | $0.050 \pm 0.052$ | $0.005 \pm 0.01$ | 38 | 1990 | $0.054 \pm 0.047$ | $0.005 \pm 0.009$ | 42 | 1684 |

<sup>§</sup> $\mu\text{DoC}$ : mean depth of coverage.
<sup>†</sup> CV: coefficient of variation of coverage.
<sup>‡</sup> $\mu\text{Ho}$ : mean observed heterozygosity across all SNPs within a given gene.

We further validated these results by comparing the observed heterozygosity (Ho) of gCNVs with that of genes classified as single-copy using the haploid dataset. Heterozygous positions in haploid DNA are not expected except for sequencing errors or when reads from multi-copy regions align at the same position in the reference genome, generating pseudo-heterozygotes. As expected, the average Ho of the haploid DNA (µHo = 0.028 ± 0.07) was much lower than that of the diploid DNA of the Swedish cline dataset (µHo = 0.246 ± 0.14) and the *P. abies-P. obovata* dataset (µHo = 0.123 ± 0.13). Note that the lower Ho in the *P. abies-P. obovata* dataset compared to the Swedish cline dataset is probably due to a Whalund effect, as the population structure is much stronger. More importantly, for both dataset, genes classified as gCNVs using the diploid data have a much higher Ho in the haploid data than those classified as single-copy (Welch Two Sample *t*-test, all *p*-value < 2.2e^-16^, Fig. 2C and Table 1). Taken together, our results support a conservative but robust detection of gCNVs from short-read exome capture data, although we can’t rule out a small proportion of false positives.

### gCNVs are widespread across the genome and involved in responses to biotic and abiotic factors

Across both datasets, at least 10.8% of the targeted protein-coding genes of the Norway spruce and Siberian spruce show copy number variations. Using the consensus genetic map of Bernhardsson et al. (2019), we show that they are distributed across the 12 linkage groups and that some form clusters in the same genomic regions (Fig. S1). These clusters likely indicate long segmental duplications, where the same mutation event affect several genes; the highly fragmented nature of the reference genome makes it difficult to properly estimate the length of the amplicons. Gene Ontology (GO, biological process only) terms associated with the gCNVs identified in each dataset were mostly shared across the two dataset (Swedish cline dataset, 962 GO terms, *P. abies-P. obovata* dataset, 925 GO terms; Szymkiewicz-Simpson overlap coefficient = 0.93) and enriched for terms mainly related to immune response and signal transduction (Fisher’s exact test, FDR adjusted *p*-value < 0.05, Tables S1 and S2 and Fig. 2D & S2A). In addition to response to biotic factors, GO terms were also enriched (Fisher’s exact test *p*-value < 0.05 but FDR adjusted *p*-value > 0.05) for terms related to abiotic responses (e.g., response to water deprivation, cold, UV, red/far red light, and metal ions), growth and morphogenesis regulation (e.g., response to auxin and salicylic acid, cell death), and metabolic pathways (Tables S1 and S2 and Fig. S3 & S4).

### gCNVs follow the main population genetic structure but show a different pattern of diversity than SNPs

We then used principal component analyses (PCA) to explore population genetic structure obtained from SNPs and gCNVs. For gCNVs, we used the normalized depth of coverage of CNV-probes as a proxy for copy number. For SNPs the population structure was the same as in previous studies (L. Li et al. 2022; Q. Zhou et al. 2024). For the *P. abies-P. obovata* dataset, the SNP-based PCA captures the gradual change from *P. abies* genetic background to *P. obovata* genetic background, and the first two axes are strongly correlated with longitude and latitude (Fig. 3A & S5A). For the Swedish cline dataset, the population structure captured by the first two axes correlates with latitude (Fig. 3G & S5B). The population genetic structures retrieved with gCNVs were similar to that obtained with SNPs, but at a much lower resolution (Fig. 3B & H, Fig. S5C & D). The gCNV-based population genetic structure predominantly followed longitude for the *P. abies-P. obovata* dataset and latitude for the Swedish cline dataset (Fig. S5C & D). For both the datasets, the top principal components obtained from SNPs and CNV-probes were highly correlated (Fig. 3D & J). To further assess the reliability of our classification between CNV-probes and single-copy probes we carried out PCAs based on the normalized depth of coverage of single-copy probes only. The population structure was barely discernible for the *P. abies-P. obovata* dataset and showed much weaker correlation with latitude or longitude, and it was completely absent for the Swedish cline dataset (Fig. 3C & I, Fig. S5E & F). It confirms both the robustness of the classification, and that the observed population structure from the normalized DoC of CNV-probes is not a spurious signal from batch effect or library size.

**Figure 3.**
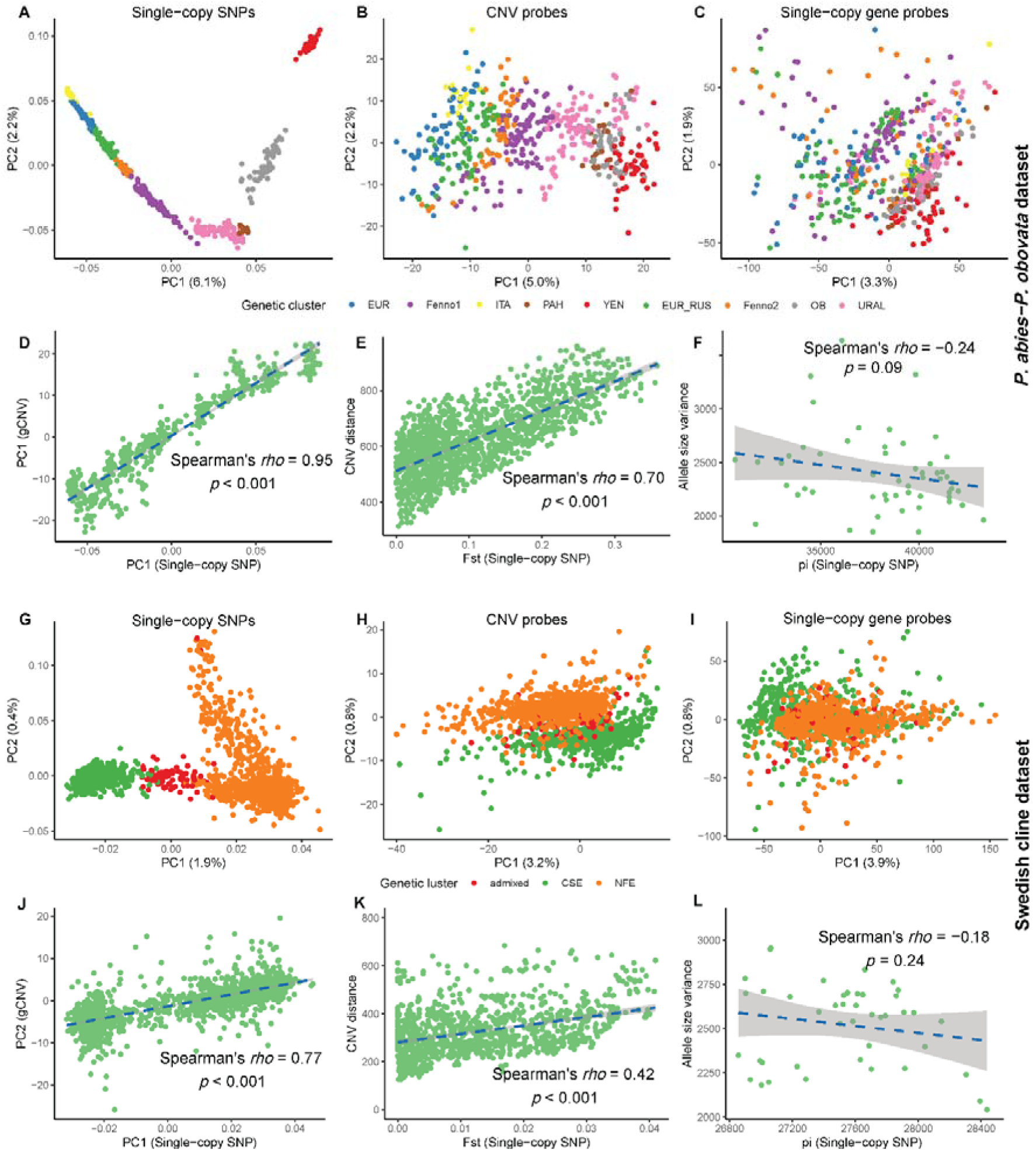
Population structure and genetic diversity obtained from SNPs and gCNVs. **A**–**C** & **G**–**I**, Principal component analyses based on single-copy SNPs, DoC of CNV-probes, and DoC of single-copy probes for the *P. abies–P. obovata* dataset (A-C) and the Swedish cline dataset (G-I). Colors correspond to genetic cluster defined by Q. Zhou et al. 2024 for the *P. abies-P. obovata* dataset and L. Li et al. 2022 for the Swedish cline dataset, respectively; **D, E and F** represent correlations between PCs scores computed using gCNVs or SNPs, population pairwise *CNV_dist_* and *F_ST_* and allele size variance (computed from gCNVs) and nucleotide diversity (summed _π_ across all neutral single-copy SNPs for each dataset), respectively for the *P. abies-P. obovata* dataset; **J,K and L,** same as D,E and F but for the Swedish cline dataset.

Estimates of Wright fixation index computed from SNPs (*F_ST_*) and gCNVs (*CNV_dist_*) were strongly correlated for both datasets (Fig. 3E & K). We then investigated the pattern of isolation-by-distance (IBD) obtained from SNPs or gCNVs. For both datasets and both types of markers, the patterns of IBD were strong, although more pronounced for SNPs (Mantel’s statistic *r F_ST_* – distance: 0.92 and 0.76, for the *P. abies-P. obovata* dataset and the *Swedish cline* dataset, respectively, all *p*-values < 1e^-4^. Fig. S6A & B; Mantel’s statistic *r CNV_dist_* - distance: 0.64 and 0.48, for the *P. abies-P. obovata* dataset and the *Swedish cline* dataset, respectively, all *p*-values < 1e^-4^. Fig. S6C & D). The strong IBD patterns as well as the strong correlations between *F_ST_* and *CNV_dist_* mean that physically and genetically close populations tend to have more similar gCNVs profiles. Taken together with the PCA, our results suggest that gCNVs globally segregate in the populations as SNPs do. However, within-population genetic diversity estimated from putatively neutral SNPs or gCNVs showed weak but negative correlations (Fig. 3F & L).

### gCNV plays a role in local adaptation and are associated to the variation of phenotypic traits

To test if gCNVs are involved in adaptation along environmental gradients, we explored the quantitative the relationship between gCNV and 19 bioclimatic variables at tree sampling locations downloaded from the Chelsa database (Fig. S7A, S8A & Table S3). First, for both datasets, we observed strong correlations between top principal components of PCAs based on normalized DoC of gCNVs or the records for the 19 bioclimatic variables (Fig. 4A & 4B). Using the same set of bioclimatic variables, we then computed pairwise population environment distances (Euclidean) to investigate the pattern of Isolation-by-Environment (IBE). Given our sampling range, we expect the environmental distance to increase along with the geodesic distance between populations. To control for it, we first regressed the *CNV_dist_* over the geodesic distances (same model as used to calculate IBD) and then tested the significance of the correlation between the residuals of this regression and the environmental distance. For both dataset we detected a significant IBE pattern for gCNVs further supporting a role for gCNVs in adaptation along environmental gradients (Spearman’s rank correlation coefficients *rho* = 0.13, *p*-value < 1e^-6^ and *rho* = 0.10, *p*-value < 1e^-6^, for *P. abies-P. obovata* and *Swedish cline* datasets, respectively).

**Figure 4.**
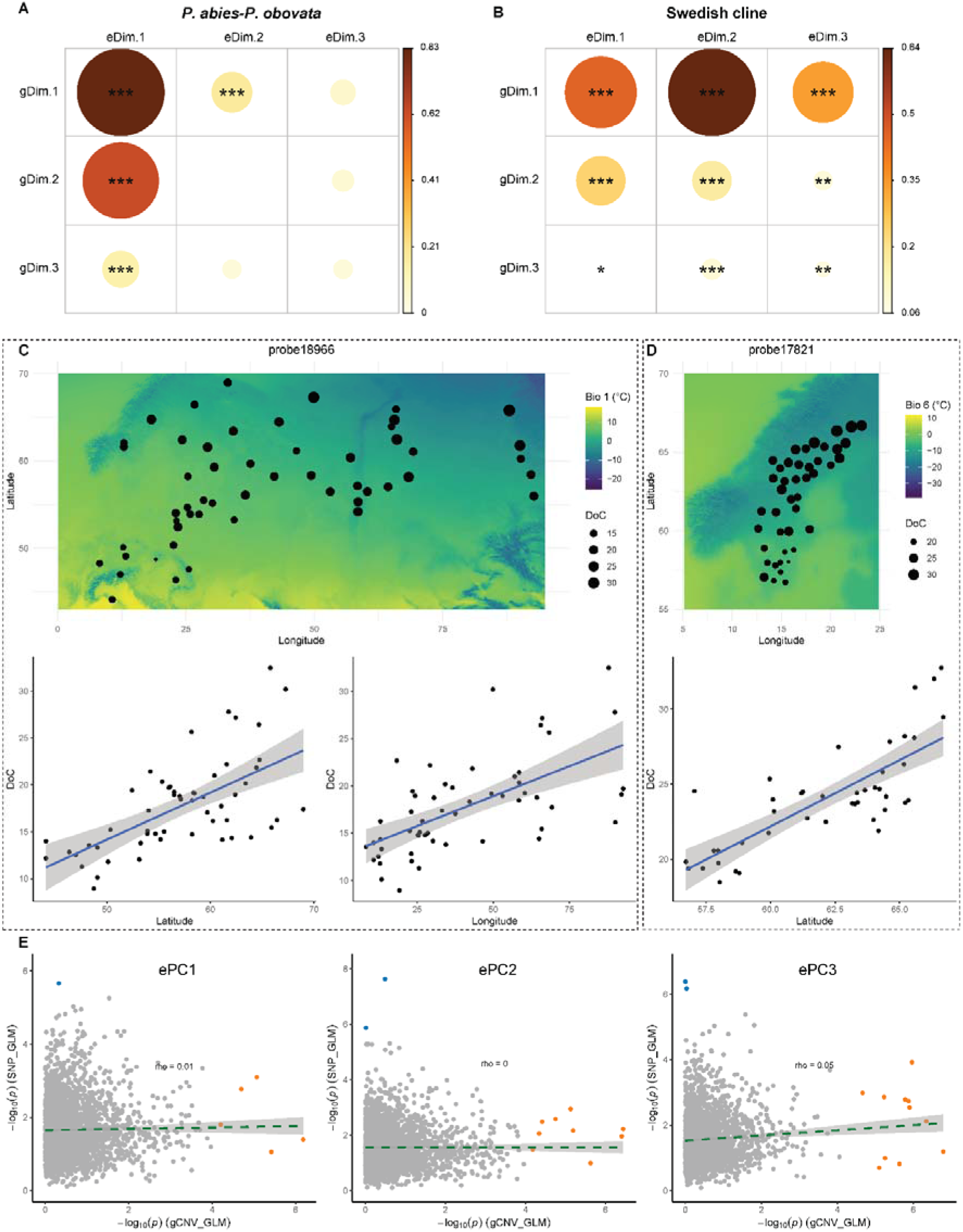
Genotype–environment associations. **A & B**, Pairwise Spearman’s correlations between the three first PCs of PCAs based on environmental data or normalized DoC of gCNVs, for the *P. abies-P.obovata* and the Swedish cline dataset, respectively. **C & D** Examples of CNV-probes that are significantly correlated with environmental variables. **C.** Top panel: Distribution of the mean normalized depth of coverage (DoC) of probe 18966 across the populations included in the *P. abies-P. obovata* dataset. The background color gradient represents the bioclimatic variable 1 (BIO1, mean annual air temperature). Bottom panel: Correlation between DoC of this probe and latitude or. **D.** Distribution of mean DoC of probe 17821 across populations included in the *Swedish cline* dataset. The background color gradient represents the bioclimatic variable 6 (BIO1, mean diurnal air temperature range). Bottom panel: Correlation between the DoC of this probe and latitude. **E,** Relationships between gCNV- and SNP-based GEA for the top three PCs of the bioclimatic variables-based PCA for the *P. abies-P. obovata* dataset. For each gene displaying CNV, the SNP with the smallest *p*-value is represented. Significant associations using the DoC of gCNVs are indicated in orange, while significant associations using SNPs are in blue. Also see Fig. S15–S18.

We then performed a series of genotype-environment association analyses using generalized linear model to detect gCNVs associated with any of the 19 bioclimatic variables or the top three PCs of a PCA based on these variables. Across all 19 variables, 56 candidate gCNVs (2.3% of all gCNVs) were identified in the *P. abies-P. obovata* dataset and 103 (3.7% of all gCNVs) in the Swedish cline dataset (Fig. S9 & S10, examples of candidate gCNVs were shown in Fig. 4C & 4D, Table S4) with only two genes identified as candidate in both datasets. One of them is *Aspartic Protease in Guard Cell 1-like* (ASPG1-like) and encodes for an aspartic protease which plays a role in drought avoidance trough abscisic acid signaling pathway; the functions of the other genes are unknown. For the top three environmental PCs, we detected 18 and 9 candidate gCNVs, respectively for the two datasets (Fig. S7 & S8); all but two overlap with the candidate gCNVs associated with individual bioclimatic variables. The low overlap between the lists of candidate gCNVs between the *P. abies-P. obovata* and the Swedish cline dataset could be explained by the different geographical range of the two datasets, different climatic drivers likely shaping their diversity. This is supported by the fact that many associations with precipitation related variables are found in the *P. abies-P. obovata* dataset where almost none were found in the Swedish dataset (Table S4). Despite the low overlap in candidate gCNVs, but as expected for genes involved in adaptation along environmental gradients, both lists were enriched for GO terms mainly associated with responses to abiotic factors and organ development (Fig. 2E, S2B, S11 & S12, Table S5 & S6). We also used a similar approach to that used by J. Chen et al. (2014) to identify gCNVs associated with latitude, using a subset of the *P. abies-P. obovata* dataset. However, no overlap was found between the significant gCNVs with those detected based on the Swedish cline dataset, even though both datasets follow latitudinal gradients.

Finally, we explored the role of gCNVs in the control of phenotypic and phenology related traits known to vary across environmental gradient in Norway and Siberian spruce by conducting a common garden experiment under controlled condition in a growth chamber (Supplementary material 1). As expected, all traits varied significantly across populations and quantitatively along latitude and longitude (Table S8). We then explored the pattern of gene copy number, phenotypic and bioclimatic variations using PCAs (Fig. 5A and Fig. S13). Top principal components of the various PCAs showed strong quantitative relationships and between each other (Fig. 5A), evidencing that gCNVs contribute to the strong local adaptation pattern through the control of phenotypic traits.

**Figure 5.**
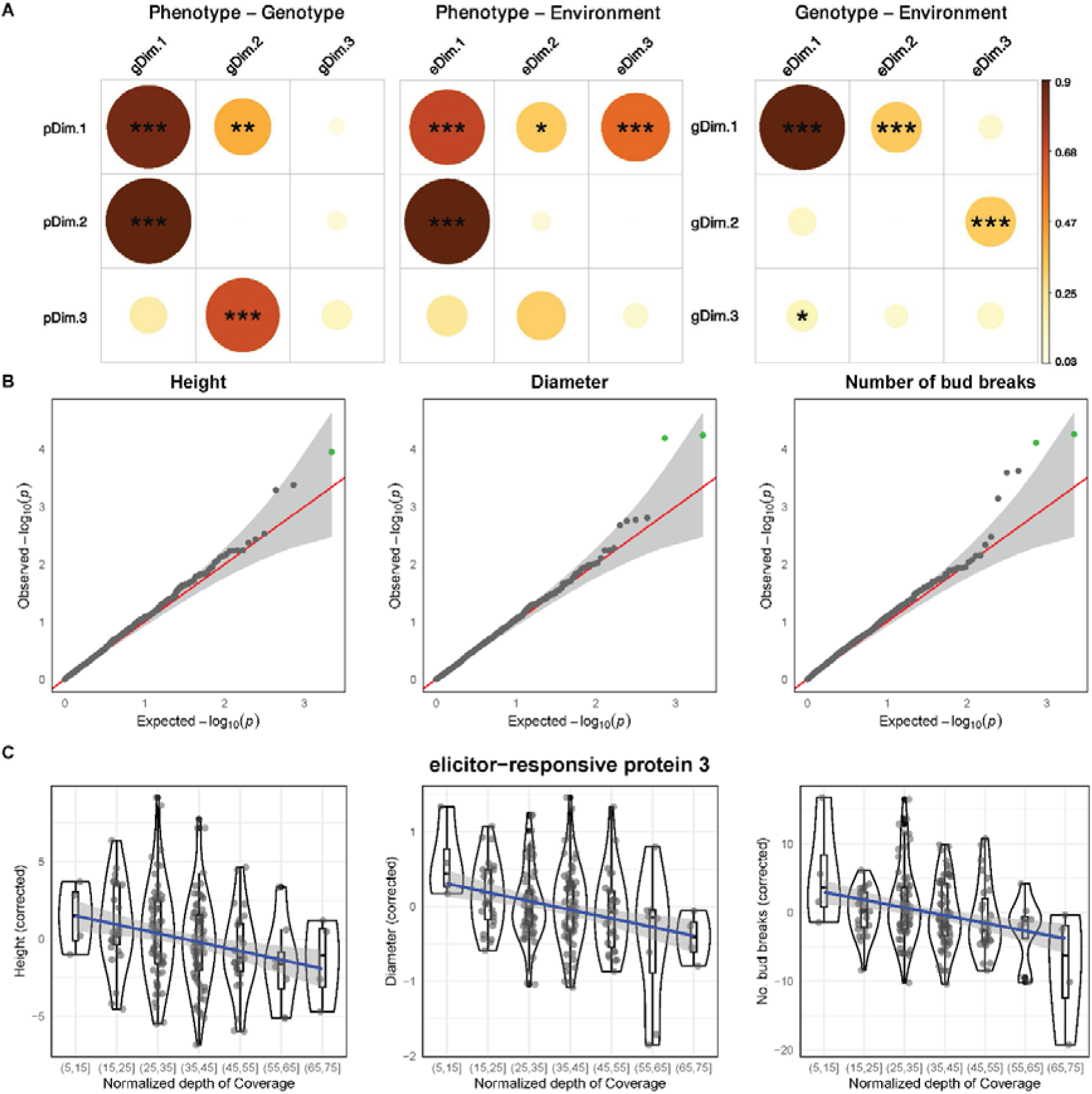
Pattern of variation of trees genotype, phenotype and original environment. **A**, Pairwise Spearman’s *rho* correlation coefficient between the top PCs of PCAs based on phenotypic data (pDim), gCNVS normalized DoC (gDim), or bioclimatic variables at tree sampling locations (eDim). Disc diameter and color scale are proportional to Spearman’s *rho* with asterisks representing the significance level (*, *p* < 0.05; **, *p* < 0.01 and ***, *p* < 0.001). **B**, Q–Q plots for genome-wide associations between gCNVs normalized DoC and three phenotypic and phenology related traits. Grey shading indicates the 95% confidence interval under the null expectation (red line), and significant associations (adjusted *p*-value < 0.2) are in green. **C**, Relationship between normalized depth of coverage and trait values (corrected for the block effect) for the candidate gene *elicitor-responsive protein 3*.

We then conducted gCNV-based GWAS using linear and generalized linear mixed models to detect candidate gCNVs involved in the control of the phenotypic traits while correcting for population structure and population of origin. Despite the limited number of samples and a strong confounding effect of population structure (Fig. S14) normalized DoC of gCNVs was significantly associated with phenotypic and phenology-related traits (tree’s height (three associations), diameter (two), number of days for bud break (one) and total number of buds (four) as well as three associations with the first PC of the PCA based of phenotypic and phenology - related data) (Fig. 5B, Fig. S14 and Tab. S4). Similar to GEA, the gCNVs that were significantly associated with the first PC were also detected as candidate genes using individual traits. Among this set of candidates, one gCNVs—annotated as *elicitor-responsive protein 3*—stands out as it seems to be involved in the control of three traits (height, diameter, and number of buds, Fig. 5C). This gene is primarily involved in the control of immune response but its over expression has been shown to reduce growth in *A. thaliana* (Jing et al. 2020; Wang et al. 2024), probably through a metabolic trade-off as often observed for amplification of resistance genes (e.g. Guillemaud et al. 1999).

### Adaptive gCNVs cannot be detected using SNPs as markers in GEA and GWAS

To test whether the same genes could have been detected using SNP data alone, we repeated the GEAs and the GWAS but using the SNPs located within the gCNVs as markers. For GEA, we used both, a population-based approach (*Bayenv2*) and an individual-based approach (same generalized linear model as for gCNVs). We also used the same linear and generalized linear models for SNP-based GWAS as for gCNVs-based GWAS. For both GEA and GWAS we used putatively neutral SNPs located within single-copy genes to control for population structure. To err on the side of caution and ensure comparability with the gCNV-based analyses, we only retained one SNP per probe and selected the one with the largest Bayes factor (*bayenv2*) or smallest *p*-value. Across all analyses only eight genes over a total of 494 significant associations (1.6%) were detected as candidate using both types of genotypes, SNPs or normalized DoC as a proxy for copy number (Tab. S4). This is further evidenced by the absence of correlation between the *p*-values obtained for GEA and GWAS using either type of genotypes (Fig. 4E and S15–S19). Whether through GEA or GWAS analyses, our study shows that the signature of selection for gCNVs cannot be detected by conventional approaches using SNPs as markers.

## Discussion

In this study, we investigated the role of gene copy number variations (gCNVs) in adaptation along environmental gradients in Norway spruce and Siberian spruce. We built a comprehensive framework for population genetic analyses with gCNVs, from their identification in exome capture sequencing data to population structure analyses and identification of candidate gCNVs contributing to local adaptation and associated with quantitative traits. We showed that gCNVs are numerous and widespread across the genomes of the two species. We globally observed the same patterns of population structure with gCNVs as with SNPs, and similar patterns of isolation by distance. A significant proportion of gCNVs were detected as candidate genes for adaptation, genes that would not have been detected using classical approaches with SNPs.

### Opportunities and limitations of using short-read exome capture data to call gCNVs

Exome capture sequencing using short reads is a cost-effective method to generate extensive population-level data. It is particularly useful for non-model species, where the reference genome is often highly fragmented, and/or for species with giga-genomes where sequencing the whole genome of many individuals is too costly. However, this sequencing technology is prone to significant variation in read depth of coverage (DoC) due to various confounding factors that make it difficult to detect copy number variations (Krumm et al. 2012). Nevertheless, exome capture targets the gene-space which is typically well annotated in reference genomes and shows a high degree of conservation (e.g. in *A. thaliana*, Igolkina et al. 2024). Thus, pseudo-SNPs are more likely to originate from gene copy number variations and paralogs than from other types of repeats, such as transposable elements. Here, we used the unsupervised machine learning-based clustering method implemented in the *rCNV* R package (Karunarathne et al. 2023) to detect pseudo-SNPs (referred to as ‘deviant’ SNPs in the *rCNV* method). As it is an indirect method that relies mainly on statistics derived from allelic DoC ratios and apparent excess of heterozygotes calculated at the SNP level it can be prone to both false positives and false negatives.

#### False positives

Various error-prone factors may influence the detection and some of the ‘deviant’ SNPs used for the classification are likely to be simply sequencing errors. As we expect these errors to be randomly distributed across the genome, we only considered probes containing at least 30% of the SNPs flagged as ‘deviant’ as candidates for multi-copy genes, using their local density as a source of information. Other false positives may arise from paralogs (e.g. fixed duplicates), as SNPs within paralogous genes may also be flagged as ‘deviant’. With exome capture data, the absolute depth of coverage is too variable from one targeted region to another to be used to identify fixed paralogs by fold change (Neves 2013; Krumm et al. 2012). However, our study has shown that the relative depth of coverage of a given locus across samples can be reliably used after correcting for confounding factors (e.g. batch effect, GC content, Fromer et al 2012). It is then possible to use a clustering approach (e.g. MCLUST, Scrucca et al. 2016) to define copy number groups and filter out CNV-probes for which the best model would support a unique group. Such probes are likely to bind to fixed or nearly fixed paralogous copies. In contrast to previous studies in conifers using whole genome data (Nystedt et al. 2013; Warren et al. 2015; Niu et al. 2022; Jang et al. 2024), only a small fraction of candidate multi-copy genes was identified as paralogs in our study. A first explanation comes from the design of the exome capture experiment, which avoids large gene families because the same probe would bind to too many loci (e.g., Neves 2013; Vidalis et al. 2018; Milesi et al. 2024). Also, reads from duplicated copies that have accumulated divergence over a long enough time are less likely to be misaligned and therefore would not be detected. Another explanation lies in the specific features of the *rCNV* method, which has been optimized for the detection of copy number variations (e.g. the use of coefficient of variation of DoC, see also discussion in Karunarathne et al. 2023). In addition, fixed substitutions between two divergent paralogous copies have a 1:1 allelic ratio for each individual; only the apparent excess of heterozygotes induced by their presence allows their discrimination. In any case, the presence of false positives is expected to introduce random noise into the depth of coverage data, which would attenuate the pattern of population structure or any quantitative relationship with the number of copies (e.g. IBD, GEA, GWAS). Our results are therefore conservative and appear robust to the potential presence of undetected false positives.

#### False negatives

False negatives would be mainly due to a lack of statistical power when classifying the SNPs with the *rCNV* method. A too low depth of coverage of a given region or too few heterozygous individuals for a given SNP would prevent a robust detection of a ‘deviant’ SNP (Karunarathne et al. 2023). Another limitation is the frequency of segregating duplicates in the dataset. Rare CNVs would result in a small number of heterozygotes with a skewed allele ratio, making their detection difficult. Similarly, the approach we used is blind to CNVs that would be strictly identical in the region captured (i.e., no pseudo-SNP present), which would also contribute to the low number of paralogs we detected. In contrast to false positives, false negatives are not expected to affect the patterns of variation observed with normalized depth of coverage for gCNVs. The various controls we have performed using normalized depth of coverage with single-copy genes also tend to indicate a rather low proportion of false negatives, or that they segregate at a low enough frequency to not generate patterns of population structure or spurious associations (Fig. 3 & S5).

#### Validation

The number of putative gene CNVs and the large sample size of our datasets would make validation of even a small proportion of gCNVs using quantitative PCR-based approaches a time-consuming and costly endeavor. Instead, we used haploid DNA from the megagametophyte, a tissue found in the seed of spruce species, as a high-throughput and cost-effective approach to validate our detection of gCNVs as, for example, in Prunier et al. (2017) and Lind et al. (2022). Expanding the probe set from 48,000 to 90,000, while targeting the same genes, increased the resolution of our validation set. It is worth noting that validation using haploid data is performed using only the density of SNPs, regardless of the depth of coverage of the different alleles. The validation step is thus performed on a different signature than the detection step, and our study demonstrates the reliability of using haploid sequencing data to validate the call of gCNVs genome-wide; given that sequencing errors are limited. In coniferous species, for example, an optimal sequencing design would involve sequencing both haploid DNA from the megagametophyte and diploid DNA from the embryo from the same seed (Bernhardsson et al. 2019; Lind et al. 2022). In addition to generating high-confidence SNP and gCNV data, such a design would also allow the phasing of the two types of polymorphism, giving access to the relative copy number carried by each homologous chromosome rather than an individual average.

### Large gCNV polymorphism in *Picea abies* and *P. obovata* populations

Over all the genes captured in our experiment (26,223) and despite conservative filtering criteria, ∼10% of them display copy number variations. This large fraction is likely to be a lower bound, as our detection approach excludes i) rare copy number variant, ii) large gene families and misses iii) copies with strictly identical sequences in the probe’s regions. Such a large number is more likely due to a high gene duplication rate than to a whole genome duplication (WGD) event followed by biased gene retention as often observed in plant species (Panchy et al. 2016). As a single WGD event occurred at the root of Pinaceae between 200 and 342 mya (Stull et al. 2021), a substantial divergence between the copies retained would be expected. Our findings are, however, consistent with recent studies in spruce species based on WGS data (Nilsson et al. 2025) or array comparative genomic hybridization (aCGH) sequencing (Prunier, Caron, Lamothe, et al. 2017; Sahli 2017). Nilsson et al. (2025) showed that since its divergence from Scots pine (*Pinus sylvestris*) ca. 130 mya, Norway spruce genome accumulated 1 Gb of genic sequences mostly through large segmental duplications, independently from transposable element activities. Sahli (2017) estimated a rate of 3*10^-5^ copy number changes per gene per generation using *Picea glauca* (white spruce) pedigrees. This estimate is within the range of gene duplication rate obtained for other eukaryote organisms (Schrider et al. 2013; Denver et al. 2009; Lipinski et al. 2011; Pan and Zhang 2007) and would explain the large polymorphism of copy number observed in the genome of Norway spruce and Siberian spruce. We identified gCNVs in all the linkage groups as well as the presence of several clusters suggesting the existence of potential gCNV hotspots (Fig. S1), as found in poplar (Prunier et al. 2019; Pinosio et al. 2016). It is also possible that these clusters, or some of them, belong to large segmental duplications encompassing several genes (e.g. Assogba et al. 2016). The use of exome capture technology however impedes us to tease apart the alternative hypotheses, but the study by Nilsson et al. (2025) would support the latter. Nonetheless, considering the spread of the gCNVs across the linkage groups most of them probably occurred from independent events, further suggesting a high gene duplication rate.

#### gCNVs and SNPs segregate in a similar way between population but may have different global fitness effect

Despite the large difference in geographical range, we found that genes showing copy number variations largely overlap between the two datasets (∼71% overlap, 2,173 genes). This is not surprising given that *P. abies* and *P. obovata* form a syngameon (Q. Zhou et al. 2024) with extensive gene flow, relatively low level of genetic differentiation (Tsuda et al. 2016; J. Chen et al. 2019; Q. Zhou et al. 2024) and a small fraction of fixed substitutions (Karunaratne et al. 2024). As other studies on different organisms previously showed (e.g. Sjödin and Jakobsson 2012; Xu et al. 2016; Cheeseman et al. 2016), we characterized a similar population structure using either the normalized depth of coverage of gCNVs or SNPs as genotypes, but with much lower resolution for the former. If this shows that gCNVs tend to segregate as SNPs between populations, it appears that there is little, if any, benefit to considering them in addition to SNPs in estimating population structure, especially since a large fraction of SNPs are likely to be effectively neutral which might not be the case for gCNVs.

In striking contrast to SNPs, we found a similar number of gCNVs in each dataset. One reason for this could be that our approach to detect gCNV is not exhaustive and is biased toward common gCNVs. On the other hand, the different patterns in numbers of SNPs and gCNVs between these two datasets may also suggest that different evolutionary forces shape the bulk of the diversity of these two types of polymorphisms, as supported by the lack of positive correlation between within population diversity of gCNVs and neutral SNPs (Fig. 3F & L). This is even more likely considering the higher estimates of the gene duplication rate compared to the point mutation rate. Several studies have suggested that gCNVs are more likely to have larger fitness effects than SNPs copies (e.g., Katju and Bergthorsson 2013; Schrider, Houle, et al. 2013; Adler et al. 2014; Langley et al. 2012; Dorant et al. 2020; Stuart et al. 2023), which likely increase with the number of copies (e.g., Adler et al. 2014; Langley et al. 2012; Katju and Bergthorsson 2013; Schrider, Houle, et al. 2013). The vast majority of gCNVs are thus expected to be highly deleterious (Schrider, Houle, et al. 2013) and should be removed by purifying selection. Conversely, high copy number CNVs that are kept in the population are more likely to have a positive fitness effect. For example, most adaptive gene amplifications conferring resistance to xenobiotics are selected against in their absence (e.g. Kondrashov 2012; Milesi et al. 2017; Weill et al. 2000; Andersson and Hughes 2009), demonstrating the intrinsic deleterious effects of such mutations. A comparison of the frequency spectrum of SNPs and gCNVs would be interesting to assess the proportion of gCNVs that are effectively neutral. It is, however, not possible with the current dataset because our detection method is biased towards common gCNVs. Also, we only access relative copy-number while absolute copy number would be needed as our study, as others before, suggests that different copy numbers may have different selective effects.

#### gCNVs are key players in adaptation along environmental gradient

The development of high-throughput sequencing technologies and the acquisition of population-level genomic data offer the opportunity to gain a comprehensive view of the genetic architecture of adaptation. This notably evidenced the role of genomic structural variations in adaptation alongside that of SNPs (e.g. Assogba et al. 2016 for large segmental duplications, Battlay et al. 2025 for inversions). Despite noisy DoC of CNVs probes, a strong confounding effect with population structure and conservative thresholds, we detected a significant number of candidate adaptive gCNVs in both datasets, indicating their role in adaptation along environmental gradient at small and large geographical scales. The low overlap in candidate gCNVs between the two datasets (only one) suggests that the variation in number of copies is likely a respond to different selective drivers in different part of the range, as illustrated by the different number of associations with precipitation-related variables found in the *P. abies-P. obovata* dataset (62 associations) and the *Swedish cline* dataset (only four, Table S4). Our study also provides support for a role of gCNVs in controlling quantitative traits involved in adaptation along the same gradients (growth traits and phenology-related traits) even though the power of the analysis was limited by a low number of individuals. This in line with Prunier et al. (2017) that showed that gCNVs are involved in the control of phenology (e.g. bud set, bud flush) and growth traits in *Picea glauca*, using parent – offspring’s trios in a breeding program. The majority of candidate gCNVs that we identified from GEA and GWAS are involved in relevant biological processes as immune response, metal ion response, flowering time regulation, and response to environmental stress and stimulus (e.g. water deprivation, cold, and far-red light). These functions were repeatedly identified in different species (Schiessl et al. 2017; Wei et al. 2023; Prunier, Caron, Lamothe, et al. 2017; Hung et al. 2024; Hardigan et al. 2016), but also in Norway and Siberian spruce using SNPs as markers (J. Chen et al. 2014; Milesi et al. 2019; L. Li et al. 2022; Z. Q. Chen et al. 2021; Karunarathne et al. 2024). The high duplication rate, multi-allelic and quantitative nature of gCNVs likely enable a dynamic and rapid response to environmental variation, whereby different copy numbers are selected along environmental gradients. On the other hand, gCNVs are likely to be more detrimental when not adaptive, and one would expect them to have a lower standing variance compared to SNPs, what could delay the response. In any case, our study shows that gCNVs, alongside SNPs, contribute to local adaptation along environmental gradients, further supporting polygenic adaptation (Milesi et al. 2019; L. Li et al. 2022; Z. Q. Chen et al. 2021).

#### The effect of gCNVs is not captured by SNPs

Last but not least, adaptive gCNVs could not have been identified using SNP data alone. On the short term, gCNVs can be selected for two main reasons: gene dosage (e.g. Kondrashov 2012) and fixation of a permanent heterozygote advantage (segregation avoidance model, Milesi et al. 2017; Spofford 1969). Under the dosage scenario, the adaptive potential is independent of the observed allele frequencies, whereas under the segregation avoidance model, the SNPs trapped in the duplication would show an apparent excess of heterozygotes (Lenormand et al. 1998) and would have probably been filtered out of the dataset using standard SNP filtering procedure. Although most of the genome scans, GWAS and GEA methods were primarily developed for analyzing SNP data, they have proved effective in detecting adaptive structural variations (SVs) using shift in SNPs frequencies as markers. If this is particularly true for balanced SVs such as inversions (Ayala et al. 2019; Kennington et al. 2007; Koch et al. 2021; Westram et al. 2023), it is much less so for unbalanced mutations, as deletions or copy number variations (e.g. Mérot et al. 2020). For example, Redon et al. (2006) found that causal CNVs in humans were poorly predicted by neighboring SNPs. Prunier, Caron, Lamothe, et al. (2017) found no overlap between gCNVs and SNPs in *P. glauca* and Harris et al. (2024) obtained much higher polygenic scores for various quantitative traits when including CNVs in addition to SNPs. gCNVs can thus be considered largely untapped genetic variation that may even explain a fraction of phenotypic and / or fitness variation, and, to some extent, part of the ‘missing-heritability’ (Maher 2008), as for other structural variants (e.g., Yao Zhou et al. 2022). Since SNPs do not capture their effects, a comprehensive understanding of their role at short evolutionary timescales requires characterizing gCNVs *a priori* and including them in genomic studies in a more systematic way, rather than to investigating them *a posteriori* from already identified candidate genes. Such information would also be particularly relevant in quantitative genomics, for instance in the context of plant and animal breeding.

### Conclusion

The last decade has seen a giant leap in the availability of high-throughput sequencing data. These data have primarily been used to analyze SNP variation across populations. However, our study has shown that short-read exome capture data can be reliably used for studying gene copy number variations. The use of old data and the recognition of gCNVs as an additional type of polymorphism opens the door to additional studies to fully capture their role on short evolutionary timescales. Our study also highlights the importance of incorporating gene CNVs into population and quantitative genomic studies to address important questions in evolutionary biology. This would require development/adaptation of current analytical methods, and our study has taken a step in this direction. Whether the pattern we observed in Norway spruce and Siberian spruce is restricted to species with similar biological characteristics (i.e., large distribution gradient, extensive gene flow, inability to track ecological niche shift) or if it is a more global pattern remains an open question. In any case, our study is an incentive to systematize the study of gene CNVs.

## Material and Methods

### Sampling and sequencing

This study combined previously published genomic datasets for approximately 2300 individual trees from different studies, but sequenced using the same exome capture experiment (Illumina sequencing of pair-end short-reads with a set of 40,018 probes targeting 26,219 genes, Vidalis et al. 2018). We defined two datasets at different geographical scales (Fig. 1):

- First, the “*P. abies-P. obovata*” dataset consists of 542 spruce individuals sampled in 55 populations along a longitudinal gradient from Western Europe and Fennoscandia (Norway spruce) to the Yenisei River (Siberian spruce) and across a large hybrid zone between the two species (Supplementary material 1, Karunarathne et al. 2024; Q. Zhou et al. 2024).
- Second, the “*Swedish cline*” dataset is composed of 1758 individual trees collected in natural populations along a latitudinal gradient whose progeny were planted in breeding trials established by Skogforsk (The Forestry Research Institute of Sweden, Supplementary material 2, L. Li et al. 2022). For population-based analysis, we gathered geographically close-by individual trees belonging to a same genetic cluster into a same population following Li et al. (2022).

In addition, we extracted haploid DNA from the megagametophyte, a haploid tissue found in the seed, for an additional 180 individuals from 36 populations covering a comparable geographic range to the *P. abies-P. obovata* dataset to capture the same genetic diversity (Fig. 1, Supplementary Material 3). We used haploid DNA as a high-throughput and time-efficient approach to cross-validate the list of candidate gene CNVs established from the diploid DNA (see below). To isolate the megagametophyte, we dissected the seeds according to the protocol of García and Escribano-Ávila (2016). We used the Nucleospin PLANT II minikit (Macherey-Nagel GmbH & Co. KG, Germany) to extract haploid DNA from megagametophytes. The haploid DNA was then sequenced by the same company as the diploid DNA (RAPiD Genomic, USA), using the same exome capture sequencing technology, but with an expanded probe set: we targeted the same genes as for the diploid DNA, but increased the number of probes per gene by designing an additional set of 51,982 probes (92,000 in total). The raw-reads were deposited in NCBI (https://www.ncbi.nlm.nih.gov/sra, BioProject PRJNA1261476).

### DNA mapping and SNP calling

For diploid DNA, we downloaded the raw-reads from NCBI (https://www.ncbi.nlm.nih.gov/sra, BioProject PRJNA511374, PRJNA731384, and PRJNA1007582). We then followed the same pipeline as used in Q. Zhou et al. 2024 to perform quality control of the raw-reads, sequence alignment against the *P. abies* reference genome (v1.0, Nystedt et al. 2013), SNP calling and filtering. Briefly, after mapping (BWA-MEM v 0.7.17, H. Li et al. 2009), PCR duplicates were removed using PICARD v 2.27.4 (https://github.com/broadinstitute/picard/), followed by genotype identification carried out with GATK HaplotypeCaller (v 4.1.4.1) individually and SNP calling using GenotypeGVCFs across all samples jointly. Hard-filtering was performed to filter out SNPs of low quality using the following criteria: QD < 2.0, MQ < 40.0, SOR > 3.0, QUAL < 20.0, MQRankSum < -12.5, ReadPosRankSum < -8.0, and FS > 60.0. The final datasets contained 254,360 and 758,429 SNPs for the *Swedish cline* and *P. abies-P. obovata* dataset, respectively, and included both putatively neutral and non-neutral SNPs. Only putatively neutral SNPs (i.e. SNPs located in intron or intergenic regions, or being synonymous in protein coding sequences) with call rate larger than 0.7 and minor allele frequencies larger than 0.01 were kept for population structure analyses. We used exactly the same pipeline for haploid DNA; note that we kept the parameter for sample ploidy equal to 2 in GATK (--sample-ploidy 2) to allow the software to call heterozygote positions.

### Identification and quantification of gene copy number variations

To identify gene copy number variations from our exome capture data, we first identified SNPs located in putative multi-copy genomic regions. We then normalized the sequencing depth of coverage of these multi-copy regions and defined copy number genotypes. Finally, we excluded regions for which there was no variation in copy number between individuals (i.e., paralogs). The whole pipeline is detailed below and represented in Fig. 6.

**Figure 6:**
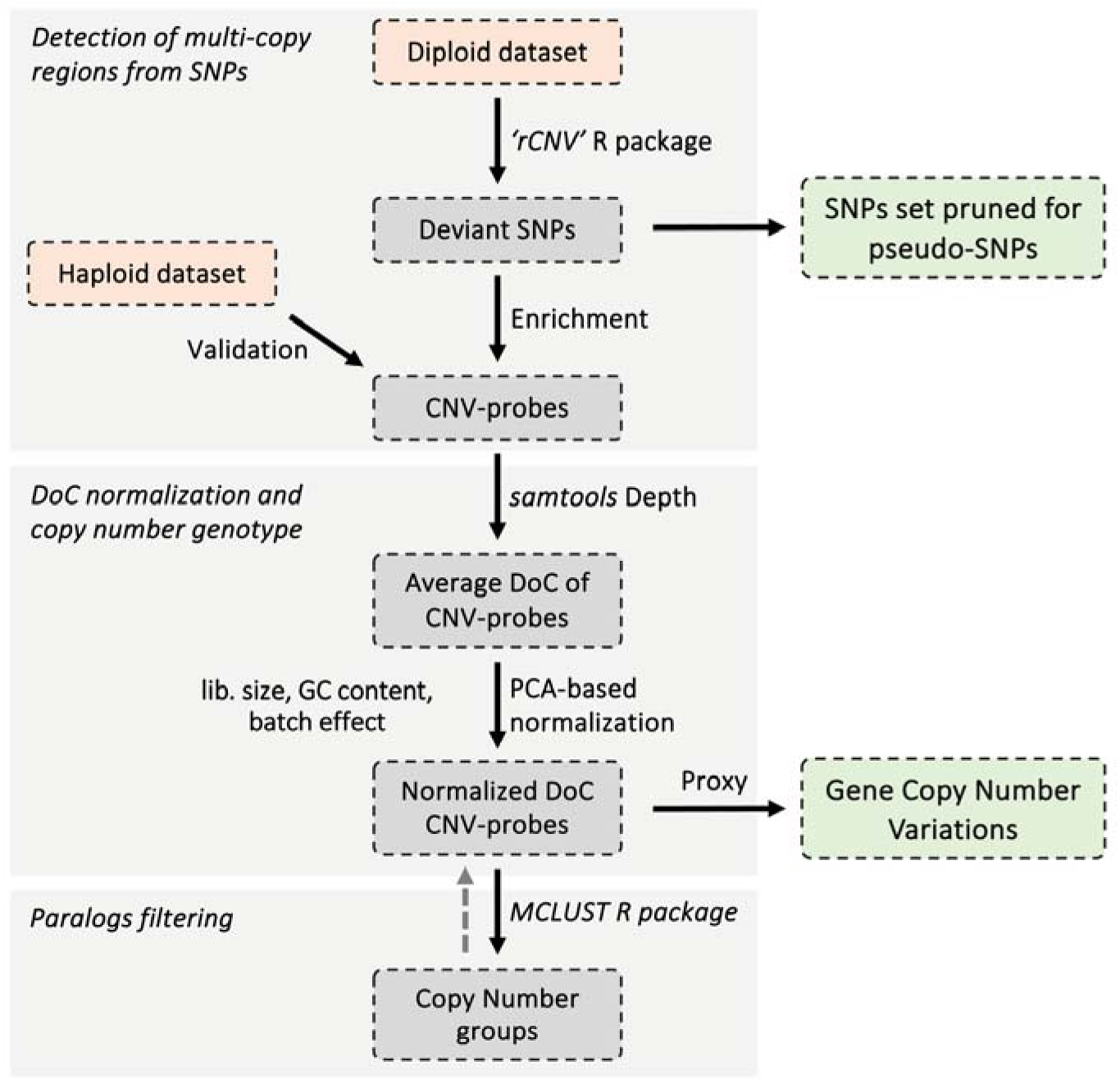
Detection and quantification of gene copy number variations from exome capture data. The main steps of the workflow are represented by the three light gray boxes and are described in detail in the Materials and Methods section. Orange boxes are input data sets, dark gray boxes are intermediate data sets, and green boxes are output data sets used for downstream analyses.

#### Detection of multi-copy regions from SNPs

As a first step, we used the approach for CNV and paralogous regions discovery from SNP data optimized for reduced genome representation sequencing data implemented in the *rCNV* R package (v1.3.9 Karunarathne et al. 2023). Briefly, it combines information from the apparent excess of heterozygotes from Hardy-Weinberg equilibrium with the deviation from the expected mean and variance of allelic depth of coverage (DoC) ratios across heterozygotes sites to classify each SNP as ‘deviant’ (i.e. putatively located in multi-copy regions) or not. We first corrected allelic DoC for genotype misclassification (where genotype call and allelic DoC do not match) and odd-numbered DoC using the *ad.correct* function. Given the scale of our sampling, we controlled for population structure when calculating the apparent excess of heterozygotes (using the *Fis* parameter in the *allele.info* function). As exome capture can introduce a bias in depth of coverage towards the reference allele (Lelieveld et al. 2015), we followed the recommendations of Karunarathne et al. (2023) and used the average allele DoC ratio across all individuals (*p.all*) as the expected allelic DoC ratio in heterozygotes (instead of *p* = 0.5). We then used the unsupervised *K-means* clustering approach to classify the SNPs as “deviant” or not (*cnv* function, *filter=kmeans* parameter). A significant proportion of false positives and false negatives are inevitable when calling CNVs from SNP data (Karanurathne et al., 2023). Therefore, in a second step, we used the density of “deviant” SNPs in each target region to further refine the list of candidate gene CNVs. Each probe used for the exome capture is 120 bp long, and we defined the targeted region of each probe by extending the sequence of the probe by 100 bp on both sides, resulting a total of 320 bp region per probe. Probes with a number of SNPs greater than two and a proportion of SNPs classified as “deviant” greater than 0.3 within the 320 bp targeted region were classified as “CNV-probes”. All the other probes with at least two SNPs were considered as “single-copy probes”. Genes which contained at least one CNV-probe were considered multi-copy genes. Conversely, only those genes which only contained probes categorized as single-copy probes were treated as single-copy genes. Using the same classification, we generated a clean SNPs dataset keeping only SNPs located within single-copy genes, resulting in 136,803 and 356,940 SNPs for the Swedish cline and the *P. abies-P. obovata* dataset, respectively.

#### Validation of multi-copy regions

We used the SNP dataset obtained from haploid DNA to validate the set of genes classified as gCNVs. As the megagametophyte is a haploid tissue, heterozygous positions are not expected unless there is a sequencing error or if reads from multiple copy regions are stacked at the same position during alignment to the reference genome. Therefore, for each gene independently, we calculated the observed heterozygosity across all the probes it contained and compared it between genes classified as single-copy or multi-copy, the latter being expected to have higher heterozygosity.

#### DoC normalization and copy number genotypes

Exome capture is a molecular hybridization-based sequencing technology, and the sequencing depth of coverage (DoC) of each position decreases with distance from the center of the probe (Neves 2013; Vidalis et al. 2018). Therefore, the DoC of a given “deviant” SNP cannot be used as a direct proxy for the number of copies of the genomic region in which it falls. Instead, we used the DoC of the 60 bp region (retrieved from the alignment files using *samtools depth* v 1.15, H. Li et al. 2009) located at the center of the probe capturing that region, as suggested by Neves (2013). Many factors can also influence the depth of coverage, either globally (e.g., library size, batch effect) or more locally (e.g., probe capture efficiency, GC content, Fromer et al. 2012). To control for the effect of these factors on DoC, we used a two-step normalization procedure to generate depth matrices in which DoC represent relative copy number genotypes (i.e., the number of copies of a given gene in a given individual). First, we excluded probes with an average DoC < 5 and normalized the DoC using the *cpm.normal* function implemented in the *rCNV* R package to account for variation in library size between samples. Second, following the approach of Krumm et al. (2012), we excluded obvious outliers using a normalization method based on principal component analysis and used singular value decomposition (SVD) to examine different factors that might influence the top five principal components (PCs, Table S8). We performed linear regressions of PC loading (for individual-based factors) or score (for probe-based factors) of each PC against the following factors: population structure (represented by PC loadings of PC1 and PC2 from PCAs based on cleaned SNP data), sequencing batch, sequencing library size, coefficient of variance of individual DoC, mean probe DoC, and probe GC content. Among the PCs examined, only those with a strong correlation with the population structure were retained, while the effect of the other confounding factors was removed from the DoC (based on the adjusted *r^2^* of the regressions, Table S8) by changing the singular values of the corresponding PCs to 0 and reconstructing normalized DoC with the new singular vectors.

#### Paralogs filtering

We used a model-based hierarchical clustering approach, implemented in the R package *MCLUST* v 6.1.1 (Scrucca et al. 2016) to further refine the candidate gCNV lists. We defined discrete copy number groups from the normalized DoC distribution of each CNV-probe across samples. Two different models were tested: an equal variance model and a spherical variable variance model. For each model, the number of components (*k*, i.e. the number of copy number groups) ranged from 1 to 10. The best model, together with the corresponding optimal *k*, was selected based on the associated Bayesian Information Criterion (BIC). We then discarded the CNV-probes for which a single copy number group was identified across all individuals (*k = 1*) or two copy number groups were identified (*k = 2*) but with fewer than five individuals in any group, similar to filtering for non-variant sites or low minor-allele count variants with SNP data. The normalized DoC of the remaining CNV-probes was then used as a proxy for gene copy number in all downstream analyses.

#### Population structure analyses

For SNP data, we used the *smartpca* function implemented in *EIGENSOFT* v 7.2.0 (Galinsky et al. 2016) with default parameters using only putatively neutral SNPs pruned for pseudo-SNPs and for linkage disequilibrium (*r^2^*> 0.5) using *plink* v2.00a3LM (Purcell et al. 2007). For gCNVs, the PCA was performed on the normalized DoC of CNV-probes using the *prcomp* function (R *base* package *v 4.2.2*).

#### Population differentiation, Isolation-by-Distance, and Isolation-by-Environment

To measure genetic diversity within population, we used VCFtools v0.1.17 to calculate per-site nucleotide diversity (Nei’s _π_, Nei and Tajima 1981) for SNP data. For gCNVs, we used the allele size variance (Valdes et al. 1993) for measuring within population diversity while using normalized DoC of CNV-probes as a proxy for gene copy number:

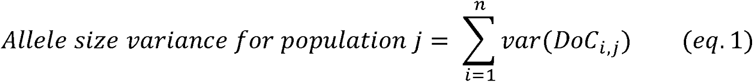

where *n* is the total number of CNV-probes, *var(Doc_i,j_)* represents the sample variance of normalized DoC for probe *i* in population *j*.

We estimated population pairwise *F_ST_* (Weir and Cockerham 1984) using VCFtools v0.1.17 (Danecek et al. 2011) based on SNP data. To ensure comparability between SNPs and gCNVs, we instead calculated population pairwise weighted Manhattan distances based on the normalized DoC of CNV-probes (hereafter *CNV_dist_*) for measuring population dissimilarity.

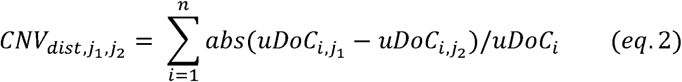

where *j*_l_ and *j*_2_ are the two populations to measure gCNV-based genetic dissimilarity, *uDoc_i_* indicates the mean depth of coverage of probe *i* across all samples.

We estimated population pairwise geodesic distances using the R package *geodist* v0.0.8 (Padgham 2021). We quantified population pairwise environmental distances using records for 19 bioclimatic variables at the population location downloaded from the *Chelsa* database v 2.1 (http://chelsa-climate.org, Table S2). The bioclimatic variables were scaled before calculating Euclidean distances between population pairs using the *dist* function in the R package *stats* v 4.2.1. We examined patterns of Isolation by Distance (IBD) by regressing either *F_ST_* or *CNV_dist_* over geodesic distances, respectively. As the environmental distance is expected to be strongly correlated with the geodesic distance between populations, we tested the patterns of Isolation by Environment (IBE) by regressing residuals of IBD over the environmental distance.

#### Testing for local adaptation

For the two main datasets, we conducted genotype-environment-association (GEA) analyses to identify candidate gCNVs involved in adaptation along environmental gradient. We used the following generalized linear model for GEA analyses controlling for population structure:

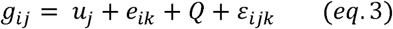

where *g_ij_* is the genotype at gCNV locus *j* of individuals *i*, *u_j_* is the mean copy number of gCNV locus *j* across all individuals, using DoC as proxy, e_ik_ is the value of environmental variable *k* at sampling location of individual *i* and *ε_ijk_* represent residuals. *Q* is a matrix containing the top PCs coordinates from a PCA based on putatively neutral SNPs pruned for LD and CNVs. The number of PCs used for controlling for population structure were determined manually based on scree plot of variance explained, as suggested by Luu et al. (2017). We performed GEA analyses for each of the 19 bioclimatic variables (Table. S3) as well as for the top three PCs from a PCA based on the bioclimatic variables, which explained > 85% of the total variance for both datasets. Because of a strong confounding effect between population structure and environmental variation in Norway spruce (Milesi et al. 2019; L. Li et al. 2022), gCNVs with a false discovery rate (FDR) < 0.2 (R package *stats* v 4.2.1) were considered as candidates contributing to adaptation along environmental gradients.

To test whether the gCNVs candidate for local adaptation could have been detected from the SNPs data alone, we also performed GEA analyses using all SNPs found within all gCNVs. Two approaches were used, both controlling for population structure. On one hand, as we did for CNVs, we used a similar population based generalized linear model by replacing the response variable with alternative allele frequency. Top three PC values for each population were obtained by averaging PC loading across all individuals within the population. SNPs with FDR < 0.2 were considered as significant associations. Second, we used *bayenv2* software (Günther and Coop 2013) to perform a GEA between each SNP and bioclimatic variable using a variance-covariance matrix of population allele frequencies (omega matrix) to control for population structure. The omega matrix was computed using *BayPass* v2.3 (Gautier 2015) based on 50,000 randomly sampled putatively neutral SNPs, pruned for multicopy regions and high linkage disequilibrium. SNPs with a Bayes factor > 20 (strong to decisive evidence according to Jeffrey’s scale, Jeffreys 1998) were considered as candidate SNPs. To be more conservative, we only retained the smallest FDR and the largest Bayes factor among SNPs within a given probe.

### Growth chamber experiment and gCNV-based genome-wide association study

A total of 230 individuals from 22 populations of the *P. abies-P. obovata* dataset (Fig. 1, Supplementary material 1) were also grown into a growth chamber in a common garden setting following J. Chen et al. (2012). Seeds were germinated and individual seedlings were grown in 8×8 cm pots randomly spread over 26 trays. The trays were spread over three benches in a growth chamber and their respective positions within and between benches were regularly randomly changed. The seedlings were first grown under continuous light (250 _μ_mol m^−2^ sec^−1^ light, 700 nm wavelength) at a temperature of 20°C for eight weeks. Then, seedlings were grown under increasing night length, where each photoperiod lasted one week, and the dark period was extended by 1.5 hr each week until reaching full dormancy in the growth chamber. The trees were kept under dormancy for a six weeks period at 12°C. Two additional growing seasons of 26 weeks were induced with a day length regime following that of 56°N (15th April to 15th of October) and a constant temperature of 20°C, each separated by a dormancy period of six weeks at 12°C adapted from Liepe et al. (2016). During the entire experiment, the substrate was kept moist by watering the seedlings regularly. Before the last growing season, we recorded the number of days before budbreak occurred and after the last growing season we measured individual height, diameter and recorded the number of branches and total number of buds per individuals.

First, we assessed patterns of local adaptation by analyzing phenotypic variation between populations using the following model:

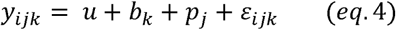

where *i*_ijk_ corresponds to the value of a given phenotypic trait measured for individual *i* from population *j* in block *k*, *ε_ijk_* represents the residuals*. b* and *p* are fixed-effects corresponding to the tray in the growth chamber and the population of origin, respectively. We used linear models to analyze height, diameter, and the number of branches (log-transformed and increased by one), as well as generalized linear models to analyze the number of buds (with a Poisson distribution) and the number of days before budbreak (with a gamma distribution and a log link function). Next, we examined the relationship between phenotypic trait values and the latitude, longitude, and altitude of the sampling location. Since these three factors are highly correlated, we analyzed them separately using the above-described model (*eq. 4*), replacing the population effect (*p*) with the corresponding factor.

Finally, we conducted genome wide association studies (GWAS) to identify gCNVs putatively involved in the control of these traits. We used the following mixed model controlling for block effect as well as population structure:

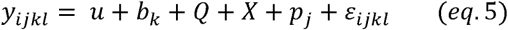

where *i_ijk_* corresponds to the value of a given phenotypic trait measured for individual *i* from population *j* in block *k*, *ε_ijk_* represents the residuals*. b* is the fixed effect corresponding to the tray. *Q* is a matrix of the first three PCs’ coordinates from a PCA based on putatively neutral SNPs from single-copy genes pruned for LD to control for population structure. *X* is a matrix of normalized DoC for CNV-probes, and *p* is the random effect corresponding to the population of origin. We used linear mixed models to analyze height, diameter, and the log-transformed number of branches increased by one. We also used generalized linear mixed models to analyze the number of buds with a Poisson distribution and the number of days before budbreak with a gamma distribution and a log link function.

#### Functional annotation and GO enrichment analyses

The protein coding sequence (CDS) of each gene included in the probe panel was extracted from the reference genome based on the generic feature annotation v1.0 (http://congenie.org/) and translated into protein sequences using *gffread* v 0.12.8 (Pertea and Pertea 2020). Functional annotation was obtained by aligning the protein sequences against the non-redundant (NR) database (https://www.ncbi.nlm.nih.gov/), the UniProtKB database (https://www.uniprot.org/), and the Pfam database (http://ftp.ebi.ac.uk/pub/databases/Pfam/releases/), using BLAST v 2.15.0+ (Johnson et al. 2008) and DIAMOND v2.1.9 software (Buchfink et al. 2021). For each protein sequence, the five most significant hits with the lowest *e*-values were selected from each database. The associated gene ontology (GO) terms (biological process only) were obtained by either ID mapping (https://www.uniprot.org/id-mapping) or InterProScan (https://www.ebi.ac.uk/interpro/search/sequence/). Functional enrichment analyses were performed with *clusterProfiler* v 4.6.2 (Wu et al. 2021) and visualized with *rrvgo* v 1.1.0 (Sayols 2023).

## Author contribution

PM and QZ conceived and designed the study, performed the experiments, analyzed the data, and wrote the manuscript. ML contributed data. PM and ML supervised the work. All authors participated in reviewing – editing tasks.

## Acknowledgement

We would like to thank Piyal Karunarathne for his assistance with code troubleshooting and for his helpful guidance on implementing specific functions in the *rCNV* R package and Jun Chen, Clémence Monod, Chen Chen and Piyal Karunarathne for their help with setting up the growth chamber experiment and traits measurement. We also would like to thank Vladimir Semerikov for his help with sampling most of the *P. obovata* material and Elena Nakvasina in providing us with additional plant materials. We also thank Øyvind Meland Edvardsen and the Norwegian Forest Seed Center, Arne Steffenrem and the Norwegian Institute of Bioeconomy Research, Luc E. Pâques and the INRAE-UMR BIOFORA Orléans, France, Muhidin Šeho and the Office for Forest Genetics, Teisendorf, Germany, Darius Danusevicius and the Vytautas Magnus university, Kaunas, Lithuania, Marcela van Loo and the Austrian Research Center for Forest, Giovanni Giuseppe Vendramin and Andrea Piotti and the Institute of Biosciences and BioResources, CNR, Italy, for sampling seeds in Norway spruce Natural populations. We thank the Swedish National Infrastructure for Computing (SNIC) for allocating computing and data storage resources for this project under the numbers NAISS 2024/6-389 and UPPMAX 2025/2-24. High-throughput sequencing costs for this study were supported by Nilsson-Ehle Endowments (43255) and Lundman’s Foundation for Botanical Studies grants from the Swedish Phytogeographic Society, awarded to Qiujie Zhou. This work was supported by Formas – a Swedish Research Council for Sustainable Development - through grant numbers 2016-00780 awarded to Martin Lascoux and 2024-02415 awarded to Pascal Milesi.

## Data availability and open science

The raw resequencing data of the newly sequenced haploid samples from this study have been deposited in the National Center for Biotechnology Information (NCBI) BioProject PRJNA1261476. All the scripts and intermediate data used in the analysis are archived in https://zenodo.org/records/17105753.

