## Supplemental tables 1-8 for "Gene copy number variation (gCNV) contributes to adaptation along environmental gradient"

**Table S1**. Enriched GO terms (biological process) of candidate gene CNVs in the Swedish cline dataset.

| **ID** | **Description** | **p value** | **p.adjust** |
| --- | --- | --- | --- |
| GO:0009626 | plant-type hypersensitive response | 4.94E-14 | 4.75E-11 |
| GO:0043068 | positive regulation of programmed cell death | 6.06E-13 | 2.92E-10 |
| GO:0019677 | NAD catabolic process | 1.98E-12 | 6.35E-10 |
| GO:0050832 | defense response to fungus | 6.91E-09 | 1.66E-06 |
| GO:0007165 | signal transduction | 3.25E-06 | 6.25E-04 |
| GO:0042742 | defense response to bacterium | 1.48E-05 | 2.38E-03 |
| GO:0016045 | detection of bacterium | 4.12E-05 | 5.67E-03 |
| GO:0009835 | fruit ripening | 9.21E-05 | 1.11E-02 |
| GO:0035336 | long-chain fatty-acyl-CoA metabolic process | 3.76E-04 | 3.85E-02 |
| GO:0009611 | response to wounding | 4.00E-04 | 3.85E-02 |
| GO:0009753 | response to jasmonic acid | 1.33E-03 | 1.16E-01 |
| GO:0009718 | anthocyanin-containing compound biosynthetic process | 1.72E-03 | 1.28E-01 |
| GO:0051707 | response to other organism | 1.83E-03 | 1.28E-01 |
| GO:0009737 | response to abscisic acid | 1.90E-03 | 1.28E-01 |
| GO:0010218 | response to far red light | 1.99E-03 | 1.28E-01 |
| GO:0050687 | negative regulation of defense response to virus | 2.77E-03 | 1.61E-01 |
| GO:0009615 | response to virus | 2.85E-03 | 1.61E-01 |
| GO:0090307 | mitotic spindle assembly | 4.08E-03 | 2.08E-01 |
| GO:0009820 | alkaloid metabolic process | 4.10E-03 | 2.08E-01 |
| GO:0010359 | regulation of anion channel activity | 4.68E-03 | 2.25E-01 |
| GO:0009625 | response to insect | 6.11E-03 | 2.75E-01 |
| GO:0010047 | fruit dehiscence | 6.28E-03 | 2.75E-01 |
| GO:0002764 | immune response-regulating signaling pathway | 7.18E-03 | 2.97E-01 |
| GO:0009862 | systemic acquired resistance, salicylic acid mediated signaling pathway | 7.84E-03 | 2.97E-01 |
| GO:0046686 | response to cadmium ion | 8.04E-03 | 2.97E-01 |
| GO:0030307 | positive regulation of cell growth | 8.33E-03 | 2.97E-01 |
| GO:0046688 | response to copper ion | 8.33E-03 | 2.97E-01 |
| GO:0009627 | systemic acquired resistance | 1.04E-02 | 3.31E-01 |
| GO:0006817 | phosphate ion transport | 1.06E-02 | 3.31E-01 |
| GO:0006826 | iron ion transport | 1.08E-02 | 3.31E-01 |
| GO:0008610 | lipid biosynthetic process | 1.13E-02 | 3.31E-01 |
| GO:1901601 | strigolactone biosynthetic process | 1.13E-02 | 3.31E-01 |
| GO:0071555 | cell wall organization | 1.14E-02 | 3.31E-01 |
| GO:0055046 | microgametogenesis | 1.39E-02 | 3.84E-01 |
| GO:0008361 | regulation of cell size | 1.49E-02 | 3.84E-01 |
| GO:0033617 | mitochondrial cytochrome c oxidase assembly | 1.50E-02 | 3.84E-01 |
| GO:0052545 | callose localization | 1.50E-02 | 3.84E-01 |
| GO:0009751 | response to salicylic acid | 1.52E-02 | 3.84E-01 |
| GO:0002239 | response to oomycetes | 1.56E-02 | 3.85E-01 |
| GO:0010150 | leaf senescence | 1.74E-02 | 3.85E-01 |
| GO:0009733 | response to auxin | 1.75E-02 | 3.85E-01 |
| GO:0009828 | plant-type cell wall loosening | 1.86E-02 | 3.85E-01 |
| GO:1900426 | positive regulation of defense response to bacterium | 1.87E-02 | 3.85E-01 |
| GO:0009409 | response to cold | 1.90E-02 | 3.85E-01 |
| GO:0006574 | valine catabolic process | 1.92E-02 | 3.85E-01 |
| GO:0006828 | manganese ion transport | 1.92E-02 | 3.85E-01 |
| GO:0016099 | monoterpenoid biosynthetic process | 1.92E-02 | 3.85E-01 |
| GO:0042178 | xenobiotic catabolic process | 1.92E-02 | 3.85E-01 |
| GO:0010039 | response to iron ion | 2.05E-02 | 3.93E-01 |
| GO:0090378 | seed trichome elongation | 2.05E-02 | 3.93E-01 |
| GO:0009624 | response to nematode | 2.12E-02 | 3.99E-01 |
| GO:0031348 | negative regulation of defense response | 2.19E-02 | 4.05E-01 |
| GO:0002213 | defense response to insect | 2.24E-02 | 4.07E-01 |
| GO:0009639 | response to red or far red light | 2.46E-02 | 4.15E-01 |
| GO:0009864 | induced systemic resistance, jasmonic acid mediated signaling pathway | 2.46E-02 | 4.15E-01 |
| GO:0045184 | establishment of protein localization | 2.46E-02 | 4.15E-01 |
| GO:0045490 | pectin catabolic process | 2.59E-02 | 4.15E-01 |
| GO:0006898 | receptor-mediated endocytosis | 2.64E-02 | 4.15E-01 |
| GO:0009617 | response to bacterium | 2.68E-02 | 4.15E-01 |
| GO:0010233 | phloem transport | 2.71E-02 | 4.15E-01 |
| GO:0015706 | nitrate transmembrane transport | 2.71E-02 | 4.15E-01 |
| GO:0009809 | lignin biosynthetic process | 2.71E-02 | 4.15E-01 |
| GO:0071577 | zinc ion transmembrane transport | 2.75E-02 | 4.15E-01 |
| GO:0050829 | defense response to Gram-negative bacterium | 2.76E-02 | 4.15E-01 |
| GO:0002237 | response to molecule of bacterial origin | 2.86E-02 | 4.23E-01 |
| GO:0002229 | defense response to oomycetes | 2.93E-02 | 4.27E-01 |
| GO:0090708 | specification of plant organ axis polarity | 3.07E-02 | 4.28E-01 |
| GO:2000070 | regulation of response to water deprivation | 3.07E-02 | 4.28E-01 |
| GO:0000272 | polysaccharide catabolic process | 3.07E-02 | 4.28E-01 |
| GO:0048544 | recognition of pollen | 3.19E-02 | 4.32E-01 |
| GO:0072593 | reactive oxygen species metabolic process | 3.19E-02 | 4.32E-01 |
| GO:0160073 | Casparian strip assembly | 3.49E-02 | 4.65E-01 |
| GO:0010087 | phloem or xylem histogenesis | 3.53E-02 | 4.65E-01 |
| GO:0010193 | response to ozone | 3.83E-02 | 4.98E-01 |
| GO:0009630 | gravitropism | 4.14E-02 | 5.29E-01 |
| GO:0006631 | fatty acid metabolic process | 4.21E-02 | 5.29E-01 |
| GO:0030245 | cellulose catabolic process | 4.40E-02 | 5.29E-01 |
| GO:0071219 | cellular response to molecule of bacterial origin | 0.043955 | 5.29E-01 |
| GO:0052544 | defense response by callose deposition in cell wall | 0.045026 | 5.29E-01 |
| GO:0055075 | potassium ion homeostasis | 0.045026 | 5.29E-01 |
| GO:1900057 | positive regulation of leaf senescence | 0.045026 | 5.29E-01 |
| GO:0048767 | root hair elongation | 0.045114 | 5.29E-01 |
| GO:0006636 | unsaturated fatty acid biosynthetic process | 0.049709 | 5.63E-01 |
| GO:0006749 | glutathione metabolic process | 0.049709 | 5.63E-01 |
| GO:0046274 | lignin catabolic process | 0.049709 | 5.63E-01 |

**Table S2**. Enriched GO terms (biological process) of candidate gene CNVs in the *P.abies-P.obovata* dataset.

| **ID** | **Description** | **p value** | **p.adjust** |
| --- | --- | --- | --- |
| GO:0043068 | positive regulation of programmed cell death | 2.46E-20 | 1.83E-17 |
| GO:0019677 | NAD catabolic process | 3.96E-20 | 1.83E-17 |
| GO:0009626 | plant-type hypersensitive response | 1.22E-19 | 3.76E-17 |
| GO:0007165 | signal transduction | 2.14E-11 | 4.96E-09 |
| GO:0050832 | defense response to fungus | 1.64E-10 | 3.03E-08 |
| GO:0042742 | defense response to bacterium | 1.63E-09 | 2.52E-07 |
| GO:0016045 | detection of bacterium | 3.71E-08 | 4.90E-06 |
| GO:0009835 | fruit ripening | 2.32E-05 | 2.69E-03 |
| GO:0009862 | systemic acquired resistance, salicylic acid mediated signaling pathway | 1.72E-04 | 1.77E-02 |
| GO:0009615 | response to virus | 2.89E-04 | 2.67E-02 |
| GO:0090307 | mitotic spindle assembly | 3.18E-04 | 2.67E-02 |
| GO:0010359 | regulation of anion channel activity | 4.68E-04 | 3.61E-02 |
| GO:0002764 | immune response-regulating signaling pathway | 5.53E-04 | 3.93E-02 |
| GO:0006898 | receptor-mediated endocytosis | 8.92E-04 | 5.89E-02 |
| GO:0009753 | response to jasmonic acid | 2.03E-03 | 1.19E-01 |
| GO:0009718 | anthocyanin-containing compound biosynthetic process | 2.31E-03 | 1.19E-01 |
| GO:2000031 | regulation of salicylic acid mediated signaling pathway | 2.31E-03 | 1.19E-01 |
| GO:0009567 | double fertilization forming a zygote and endosperm | 2.31E-03 | 1.19E-01 |
| GO:0009624 | response to nematode | 2.80E-03 | 1.31E-01 |
| GO:1900426 | positive regulation of defense response to bacterium | 2.82E-03 | 1.31E-01 |
| GO:0009625 | response to insect | 3.05E-03 | 1.34E-01 |
| GO:0045184 | establishment of protein localization | 3.35E-03 | 1.35E-01 |
| GO:0032940 | secretion by cell | 3.37E-03 | 1.35E-01 |
| GO:0030422 | siRNA processing | 4.49E-03 | 1.51E-01 |
| GO:0046688 | response to copper ion | 4.49E-03 | 1.51E-01 |
| GO:0090708 | specification of plant organ axis polarity | 4.49E-03 | 1.51E-01 |
| GO:0051707 | response to other organism | 4.53E-03 | 1.51E-01 |
| GO:0009611 | response to wounding | 4.63E-03 | 1.51E-01 |
| GO:0160073 | Casparian strip assembly | 4.77E-03 | 1.51E-01 |
| GO:0002229 | defense response to oomycetes | 5.00E-03 | 1.51E-01 |
| GO:0000302 | response to reactive oxygen species | 5.06E-03 | 1.51E-01 |
| GO:0009820 | alkaloid metabolic process | 6.45E-03 | 1.79E-01 |
| GO:1901601 | strigolactone biosynthetic process | 6.55E-03 | 1.79E-01 |
| GO:0006521 | regulation of cellular amino acid metabolic process | 6.59E-03 | 1.79E-01 |
| GO:0002239 | response to oomycetes | 6.97E-03 | 1.79E-01 |
| GO:0006865 | amino acid transport | 6.97E-03 | 1.79E-01 |
| GO:0007267 | cell-cell signaling | 7.62E-03 | 1.89E-01 |
| GO:0052544 | defense response by callose deposition in cell wall | 7.96E-03 | 1.89E-01 |
| GO:0055075 | potassium ion homeostasis | 7.96E-03 | 1.89E-01 |
| GO:0009787 | regulation of abscisic acid-activated signaling pathway | 9.69E-03 | 2.21E-01 |
| GO:0010310 | regulation of hydrogen peroxide metabolic process | 9.81E-03 | 2.21E-01 |
| GO:2000032 | regulation of secondary shoot formation | 1.05E-02 | 2.26E-01 |
| GO:0009809 | lignin biosynthetic process | 1.16E-02 | 2.26E-01 |
| GO:0010597 | green leaf volatile biosynthetic process | 1.16E-02 | 2.26E-01 |
| GO:0009737 | response to abscisic acid | 1.25E-02 | 2.26E-01 |
| GO:0010150 | leaf senescence | 1.27E-02 | 2.26E-01 |
| GO:0000304 | response to singlet oxygen | 1.30E-02 | 2.26E-01 |
| GO:0010039 | response to iron ion | 1.30E-02 | 2.26E-01 |
| GO:0080151 | positive regulation of salicylic acid mediated signaling pathway | 1.30E-02 | 2.26E-01 |
| GO:0010585 | glutamine secretion | 1.30E-02 | 2.26E-01 |
| GO:0016099 | monoterpenoid biosynthetic process | 1.30E-02 | 2.26E-01 |
| GO:0035278 | miRNA-mediated gene silencing by inhibition of translation | 1.30E-02 | 2.26E-01 |
| GO:0050687 | negative regulation of defense response to virus | 1.30E-02 | 2.26E-01 |
| GO:0010105 | negative regulation of ethylene-activated signaling pathway | 1.35E-02 | 2.26E-01 |
| GO:0009409 | response to cold | 1.35E-02 | 2.26E-01 |
| GO:0048544 | recognition of pollen | 1.83E-02 | 2.86E-01 |
| GO:0035987 | endodermal cell differentiation | 1.86E-02 | 2.86E-01 |
| GO:2000070 | regulation of response to water deprivation | 1.86E-02 | 2.86E-01 |
| GO:0071327 | cellular response to trehalose stimulus | 1.89E-02 | 2.86E-01 |
| GO:0071577 | zinc ion transmembrane transport | 1.89E-02 | 2.86E-01 |
| GO:0080143 | regulation of amino acid export | 1.89E-02 | 2.86E-01 |
| GO:0009414 | response to water deprivation | 2.14E-02 | 3.20E-01 |
| GO:0006826 | iron ion transport | 2.31E-02 | 3.39E-01 |
| GO:0009636 | response to toxic substance | 2.49E-02 | 3.61E-01 |
| GO:0010618 | aerenchyma formation | 2.61E-02 | 3.69E-01 |
| GO:1900057 | positive regulation of leaf senescence | 2.64E-02 | 3.69E-01 |
| GO:0006508 | proteolysis | 2.73E-02 | 3.69E-01 |
| GO:0002758 | innate immune response-activating signaling pathway | 2.82E-02 | 3.69E-01 |
| GO:0009800 | cinnamic acid biosynthetic process | 2.82E-02 | 3.69E-01 |
| GO:0042659 | regulation of cell fate specification | 2.82E-02 | 3.69E-01 |
| GO:0008219 | cell death | 2.87E-02 | 3.69E-01 |
| GO:0019048 | modulation by virus of host process | 2.87E-02 | 3.69E-01 |
| GO:0071555 | cell wall organization | 2.97E-02 | 3.76E-01 |
| GO:0009828 | plant-type cell wall loosening | 3.12E-02 | 3.90E-01 |
| GO:0032259 | methylation | 3.47E-02 | 4.23E-01 |
| GO:0090693 | plant organ senescence | 3.47E-02 | 4.23E-01 |
| GO:0002213 | defense response to insect | 3.66E-02 | 4.40E-01 |
| GO:0001666 | response to hypoxia | 0.037139 | 4.40E-01 |
| GO:2000067 | regulation of root morphogenesis | 0.042601 | 4.83E-01 |
| GO:2000377 | regulation of reactive oxygen species metabolic process | 0.042846 | 4.83E-01 |
| GO:0031408 | oxylipin biosynthetic process | 0.043981 | 4.83E-01 |
| GO:0009805 | coumarin biosynthetic process | 0.044871 | 4.83E-01 |
| GO:0031349 | positive regulation of defense response | 0.044871 | 4.83E-01 |
| GO:0035336 | long-chain fatty-acyl-CoA metabolic process | 0.044871 | 4.83E-01 |
| GO:0052545 | callose localization | 0.044871 | 4.83E-01 |
| GO:1901183 | positive regulation of camalexin biosynthetic process | 0.044871 | 0.482622 |
| GO:1902290 | positive regulation of defense response to oomycetes | 0.048005 | 0.510399 |
| GO:0009751 | response to salicylic acid | 0.04959 | 0.521262 |

**Table S3**. The 19 bioclimatic variables extracted from the Chelsa database v 2.1.

| **Variable** | **Variable full name** | **Explanation** |
| --- | --- | --- |
| bio 1 | mean annual air temperature | mean annual daily mean air temperatures averaged over 1 year |
| bio 2 | mean diurnal air temperature range | mean diurnal range of temperatures averaged over 1 year |
| bio 3 | isothermality | ratio of diurnal variation to annual variation in temperatures |
| bio 4 | mean annual air temperature | mean annual daily mean air temperatures averaged over 1 year |
| bio 5 | mean annual air temperature | mean annual daily mean air temperatures averaged over 1 year |
| bio 6 | mean diurnal air temperature range | mean diurnal range of temperatures averaged over 1 year |
| bio 7 | isothermality | ratio of diurnal variation to annual variation in temperatures |
| bio 8 | temperature seasonality | standard deviation of the monthly mean temperatures |
| bio 9 | mean daily maximum air temperature of the warmest month | The highest temperature of any monthly daily mean maximum temperature |
| bio 10 | mean daily minimum air temperature of the coldest month | The lowest temperature of any monthly daily mean maximum temperature |
| bio 11 | annual range of air temperature | The difference between the Maximum Temperature of Warmest month and the Minimum Temperature of Coldest month |
| bio 12 | mean daily mean air temperatures of the wettest quarter | The wettest quarter of the year is determined (to the nearest month) |
| bio 13 | mean daily mean air temperatures of the driest quarter | The driest quarter of the year is determined (to the nearest month) |
| bio 14 | mean daily mean air temperatures of the warmest quarter | The warmest quarter of the year is determined (to the nearest month) |
| bio 15 | mean daily mean air temperatures of the coldest quarter | The coldest quarter of the year is determined (to the nearest month) |
| bio 16 | annual precipitation amount | Accumulated precipitation amount over 1 year |
| bio 17 | precipitation amount of the wettest month | The precipitation of the wettest month. |
| bio 18 | precipitation amount of the driest month | The precipitation of the driest month. |
| bio 19 | precipitation seasonality | The Coefficient of Variation is the standard deviation of the monthly precipitation estimates expressed as a percentage of the mean of those estimates (i.e. the annual mean) |

**Table S4**. Number of significant CNV-probes in Genotype-environment association (GEA) and Genotype-phenotype association (GWAS). Numbers in parentheses indicates number of significant CNV-probes containing at least one significant SNPs for the same analysis with the same bioclimatic variable or trait or principal component. See table S3 for the explanation of the bioclimatic variables. See also Fig. S14-18.

| Genotype-environment association (GEA) | | | Genotype-phenotype association (GWAS) | |
| --- | --- | --- | --- | --- |
| Bioclimatic variable | *Swedish cline* | *P. abies-*  *P. obovata* | Trait |  |
| bio 1 | 4(0) | 23(0) | Height | 3(0) |
| bio 2 | 4(0) | 15(0) | Diameter | 2(0) |
| bio 3 | 1(0) | 12(0) | # of days for budbreak | 1(0) |
| bio 4 | 70(0) | 16(0) | # of buds | 4(0) |
| bio 5 | 1(0) | 16(0) | pPC1 | 3(0) |
| bio 6 | 51(2) | 14(0) | pPC2 | 0(0) |
| bio 7 | 56(1) | 5(0) | pPC3 | 0(0) |
| bio 8 | 0(0) | 23(0) |  |  |
| bio 9 | 5(0) | 0(0) |  |  |
| bio 10 | 0(0) | 19(1) |  |  |
| bio 11 | 40(2) | 14(0) |  |  |
| bio 12 | 0(0) | 4(0) |  |  |
| bio 13 | 0(0) | 4(0) |  |  |
| bio 14 | 0(0) | 5(0) |  |  |
| bio 15 | 7(1) | 1(0) |  |  |
| bio 16 | 1(0) | 3(0) |  |  |
| bio 17 | 0(0) | 5(0) |  |  |
| bio 18 | 0(0) | 3(1) |  |  |
| bio 19 | 1(0) | 6(0) |  |  |
| ePC1 | 5(0) | 5(0) |  |  |
| ePC2 | 1(0) | 9(0) |  |  |
| ePC3 | 3(0) | 11(0) |  |  |

**Table S5**. Enriched GO terms (biological process) of candidate adaptive gene CNVs identified with genotype-environmental association analysis in the *P. abies-P. obovata* dataset.

| **ID** | **Description** | **p value** |
| --- | --- | --- |
| GO:0007017 | microtubule-based process | 0.001136 |
| GO:0009807 | lignan biosynthetic process | 0.001721 |
| GO:0090307 | mitotic spindle assembly | 0.002175 |
| GO:0016045 | detection of bacterium | 0.00938 |
| GO:0000226 | microtubule cytoskeleton organization | 0.011887 |
| GO:0051302 | regulation of cell division | 0.012152 |
| GO:0010458 | exit from mitosis | 0.019116 |
| GO:1905428 | regulation of plant organ formation | 0.019116 |
| GO:0047496 | vesicle transport along microtubule | 0.021008 |
| GO:0080052 | response to histidine | 0.021008 |
| GO:0080053 | response to phenylalanine | 0.021008 |
| GO:0090158 | endoplasmic reticulum membrane organization | 0.021008 |
| GO:0010150 | leaf senescence | 0.022137 |
| GO:0098542 | defense response to other organism | 0.0223 |
| GO:0015749 | monosaccharide transmembrane transport | 0.022897 |
| GO:0015850 | organic hydroxy compound transport | 0.022897 |
| GO:0043201 | response to L-leucine | 0.022897 |
| GO:0046500 | S-adenosylmethionine metabolic process | 0.022897 |
| GO:0051231 | spindle elongation | 0.022897 |
| GO:1901140 | p-coumaryl alcohol transport | 0.022897 |
| GO:0006561 | proline biosynthetic process | 0.026663 |
| GO:0000278 | mitotic cell cycle | 0.027373 |
| GO:0009759 | indole glucosinolate biosynthetic process | 0.028541 |
| GO:0034497 | protein localization to phagophore assembly site | 0.028541 |
| GO:0043086 | negative regulation of catalytic activity | 0.028541 |
| GO:0048281 | inflorescence morphogenesis | 0.028541 |
| GO:0007020 | microtubule nucleation | 0.030416 |
| GO:0061025 | membrane fusion | 0.030416 |
| GO:0010021 | amylopectin biosynthetic process | 0.032287 |
| GO:0010336 | gibberellic acid homeostasis | 0.032287 |
| GO:0090693 | plant organ senescence | 0.032287 |
| GO:0009838 | abscission | 0.034154 |
| GO:0009914 | hormone transport | 0.034154 |
| GO:0070507 | regulation of microtubule cytoskeleton organization | 0.034154 |
| GO:0030155 | regulation of cell adhesion | 0.037879 |
| GO:1901653 | cellular response to peptide | 0.037879 |
| GO:0000422 | autophagy of mitochondrion | 0.041589 |
| GO:0010497 | plasmodesmata-mediated intercellular transport | 0.041589 |
| GO:0080168 | abscisic acid transport | 0.047129 |
| GO:0007163 | establishment or maintenance of cell polarity | 0.048969 |
| GO:0051781 | positive regulation of cell division | 0.048969 |
| GO:0070370 | cellular heat acclimation | 0.048969 |
| GO:0110126 | phloem loading | 0.048969 |

**Table S6**. Enriched GO terms (biological process) of candidate adaptive gene CNVs identified with genotype-environmental association analysis in the Swedish cline dataset.

| **ID** | **Description** | **p value** |
| --- | --- | --- |
| GO:1905168 | positive regulation of double-strand break repair via homologous recombination | 0.000488 |
| GO:0014003 | oligodendrocyte development | 0.001165 |
| GO:0015692 | lead ion transport | 0.001373 |
| GO:0045824 | negative regulation of innate immune response | 0.001691 |
| GO:0009617 | response to bacterium | 0.001825 |
| GO:0001964 | startle response | 0.001838 |
| GO:0001510 | RNA methylation | 0.002094 |
| GO:0009620 | response to fungus | 0.002289 |
| GO:0010022 | meristem determinacy | 0.002656 |
| GO:2000779 | regulation of double-strand break repair | 0.003615 |
| GO:0016045 | detection of bacterium | 0.00425 |
| GO:0010288 | response to lead ion | 0.004712 |
| GO:0009910 | negative regulation of flower development | 0.005256 |
| GO:0006357 | regulation of transcription by RNA polymerase II | 0.00603 |
| GO:0031348 | negative regulation of defense response | 0.00723 |
| GO:0048658 | anther wall tapetum development | 0.008282 |
| GO:0045087 | innate immune response | 0.008321 |
| GO:0010228 | vegetative to reproductive phase transition of meristem | 0.010341 |
| GO:0048573 | photoperiodism, flowering | 0.011766 |
| GO:2000028 | regulation of photoperiodism, flowering | 0.016554 |
| GO:0002229 | defense response to oomycetes | 0.017264 |
| GO:0009627 | systemic acquired resistance | 0.01799 |
| GO:0010359 | regulation of anion channel activity | 0.020207 |
| GO:1900425 | negative regulation of defense response to bacterium | 0.020207 |
| GO:0042981 | regulation of apoptotic process | 0.020958 |
| GO:0006282 | regulation of DNA repair | 0.02172 |
| GO:1901141 | regulation of lignin biosynthetic process | 0.02172 |
| GO:0009741 | response to brassinosteroid | 0.023976 |
| GO:0002237 | response to molecule of bacterial origin | 0.028025 |
| GO:0080147 | root hair cell development | 0.028221 |
| GO:0045892 | negative regulation of DNA-templated transcription | 0.029572 |
| GO:0009753 | response to jasmonic acid | 0.030766 |
| GO:0043484 | regulation of RNA splicing | 0.031727 |
| GO:0009733 | response to auxin | 0.03266 |
| GO:0048653 | anther development | 0.033542 |
| GO:0052544 | defense response by callose deposition in cell wall | 0.034465 |
| GO:0010102 | lateral root morphogenesis | 0.03729 |
| GO:0008219 | cell death | 0.038252 |
| GO:0009820 | alkaloid metabolic process | 0.040202 |
| GO:0000056 | ribosomal small subunit export from nucleus | 0.0421 |
| GO:0009435 | NAD biosynthetic process | 0.0421 |
| GO:0042754 | negative regulation of circadian rhythm | 0.0421 |
| GO:0090520 | sphingolipid mediated signaling pathway | 0.0421 |
| GO:0016540 | protein autoprocessing | 0.046213 |
| GO:0030220 | platelet formation | 0.046213 |
| GO:0045836 | positive regulation of meiotic nuclear division | 0.046213 |
| GO:1903013 | response to differentiation-inducing factor 1 | 0.046213 |
| GO:1904357 | negative regulation of telomere maintenance via telomere lengthening | 0.046213 |
| GO:2000069 | regulation of post-embryonic root development | 0.046213 |

**Table S7: Life history traits variation across geography and topography in a growth chamber experiment**

|  |  | Block | |  | Place | |  | Population | |  | Latitude | | | |  | Longitude | | | |  | Altitude | | | |  |  |
| --- | --- | --- | --- | --- | --- | --- | --- | --- | --- | --- | --- | --- | --- | --- | --- | --- | --- | --- | --- | --- | --- | --- | --- | --- | --- | --- |
| Phenotype |  | *t* | *p* |  | *t* | *p* |  | *t* | *p* |  | *b* | *se* | *t* | *p* |  | *b* | *se* | *t* | *p* |  | *b* | *se* | *t* | *p* |  | Model |
| Height |  | **2.62** | **<0.001** |  | 0.06 | 0.811 |  | **3.65** | **<0.001** |  | **-0.106** | **0.047** | **-2.27** | **0.025** |  | **-0.028** | **0.012** | **-2.30** | **0.022** |  | -0.001 | 0.001 | -0.86 | 0.392 |  | Linear model |
| Diameter |  | **3.47** | **<0.001** |  | 0.12 | 0.732 |  | **2.49** | **0.002** |  | **-0.030** | **0.008** | **-3.78** | **<0.001** |  | **-0.005** | **0.002** | **-2.27** | **0.024** |  | 0.000 | 0.000 | 1.34 | 0.182 |  | Linear model |
| # of branches |  | 1.02 | 0.439 |  | 0.25 | 0.615 |  | **7.31** | **<0.001** |  | **-0.059** | **0.007** | **-8.55** | **<0.001** |  | **-0.014** | **0.002** | **-7.78** | **<0.001** |  | **0.001** | **0.000** | **5.04** | **<0.001** |  | Linear model (log-transformed response) |
| # of buds |  | **3.84** | **<0.001** |  | 0.04 | 0.836 |  | **2.52** | **0.001** |  | **-0.025** | **0.004** | **-6.34** | **<0.001** |  | **-0.004** | **0.001** | **-3.49** | **<0.001** |  | **0.000** | **0.000** | **2.64** | **0.008** |  | Generalized linear model (Poisson) |
| # of days for  budbreak |  | 0.95 | 0.541 |  | 0.31 | 0.576 |  | **2.52** | **0.002** |  | **-0.015** | **0.004** | **-3.29** | **0.001** |  | **-0.005** | **0.001** | **-4.42** | **<0.001** |  | **0.000** | **0.000** | **2.65** | **0.009** |  | Generalized linear model  (Gamma(Link = log)) |

Number of degrees of freedom were 25, 1 and 17, respectively for the Block, the place and the population fixed-effect and 1 for each geographic or topographic variable.

Vertical doubled-dotted lines separate independent models. The “Block” fixed-effect was conserved for each model with geographic or topographic variables.

Significant effects are bolded.

**Table S8**. Correlation between the top 5 principal components (PCs) of PCA based on DOC of probes and confounding factors which might induce systematic bias into the DOC matrix. Each cell represents the adjusted r^2^ from the linear regression of each PC over corresponding factor. PCs removed from the matrix were indicated in bold. Batch effect is not applicable to the Swedish cline dataset as the samples were mainly sequenced in one project. Sample CV of DOC: coefficient of variance of DOC for each sample.

|  | Swedish cline dataset | | | | | P.abies-P.obovata dataset | | | | |
| --- | --- | --- | --- | --- | --- | --- | --- | --- | --- | --- |
|  | PC1 | **PC2** | PC3 | PC4 | PC5 | PC1 | **PC2** | **PC3** | **PC4** | PC5 |
| Population structure^$^ | 0.01 | 0.14^*^ | 0.03 | 0 | 0.01 | 0.06 | 0.27^*^ | 0.22^*^ | 0.22^*^ | 0.01 |
| Batch effect | NA | NA | NA | NA | NA | 0.64^*^ | 0.42^*^ | 0.18 | 0.13^*^ | 0.18^*^ |
| Library size | 0.61^*^ | 0.16^*^ | 0 | 0.03 | 0.01 | 0.07 | 0.48^*^ | 0.03 | 0 | 0.01 |
| Sample CV of DOC | 0.55^*^ | 0.14^*^ | 0.03 | 0.01 | 0.04 | 0.04 | 0.13^*^ | 0 | 0 | 0 |
| Probe mean DOC | 0.1 | 0.01 | 0.01 | 0 | 0.52^*^ | 0.29^*^ | 0.12^*^ | 0.07 | 0.03 | 0.03 |
| Probe GC content | 0.33^*^ | 0.08 | 0.01 | 0.02 | 0 | 0.03 | 0.21 | 0.05 | 0.02 | 0.01 |

^$^ We used the top 2 PCs from PCA results on neutral single-copy SNPs as indicator of the population structures.

^*^ p value < 1e
