## Supplemental figures 1-19 for "Gene copy number variation (gCNV) contributes to adaptation along environmental gradient"

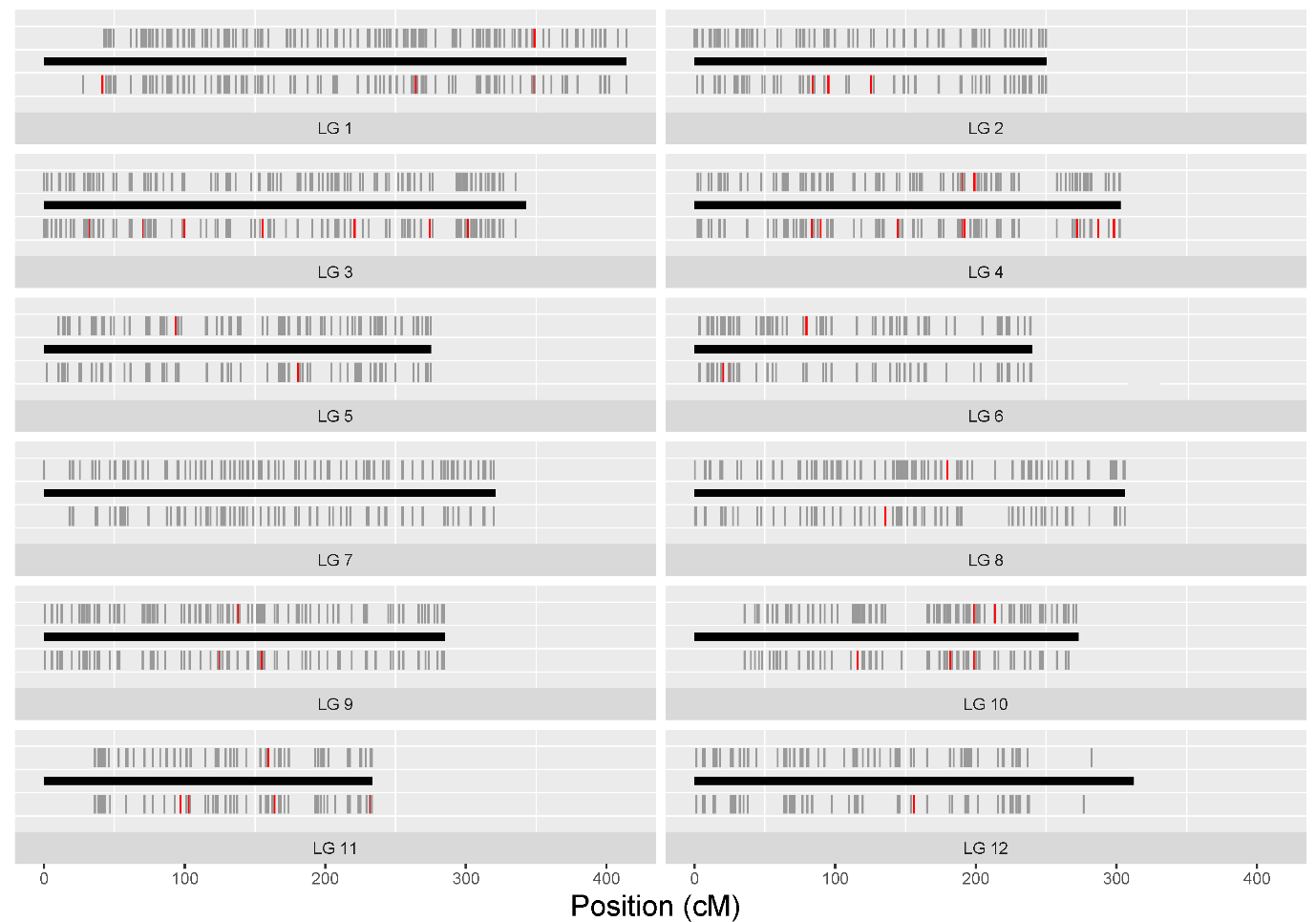
**Figure S1.** gCNV distribution across the 12 linkage groups (LG) of spruce genome. The black horizontal bar indicates the length of the LG in centiMorgan (cM). Candidate adaptive gCNVs, identified using genotype-environment association analyses, are indicated by red bars. For each linkage group, genes displaying CNVs in the *Swedish cline* dataset are represented above the black bar and below the bar are genes displaying CNVs in the *P. abies-P. obovata* dataset. A total of 22413 out of 40012 probes were assigned to the 12 LGs according to the genetic map developed by Bernhardsson et al. (2019).

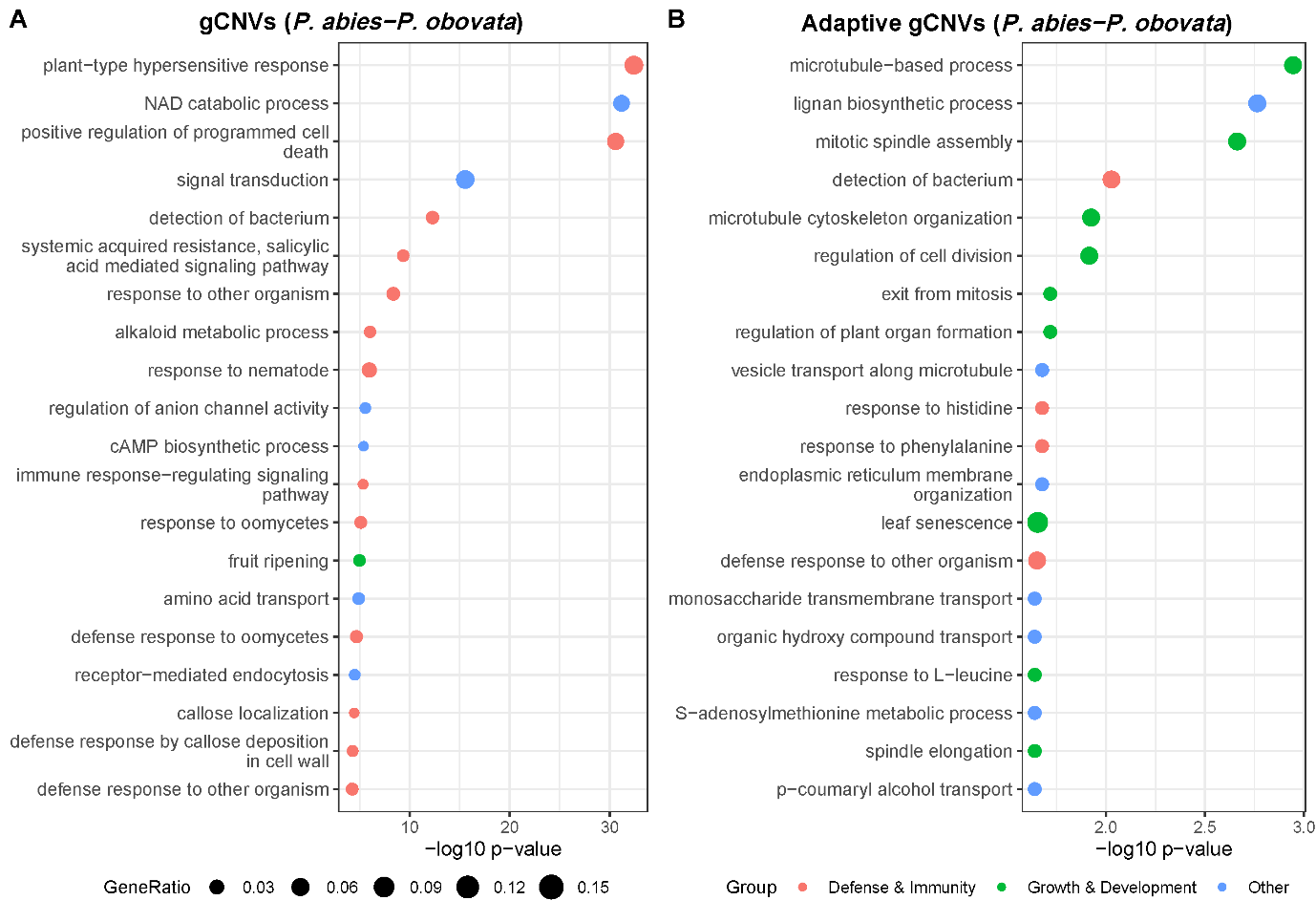

**Figure S2.** Gene Ontology (GO, biological processes) enrichment analysis of gCNVs (**A**) and candidate adaptive gCNVs detected using GEA (**B)** for the *P. abies-P. obovata* dataset. Dot size is proportional to gene ratio. Only top 20 GO terms with a *p*-value < 0.05 are shown. See also Tables S4 and S5 for full list of enriched GO terms.

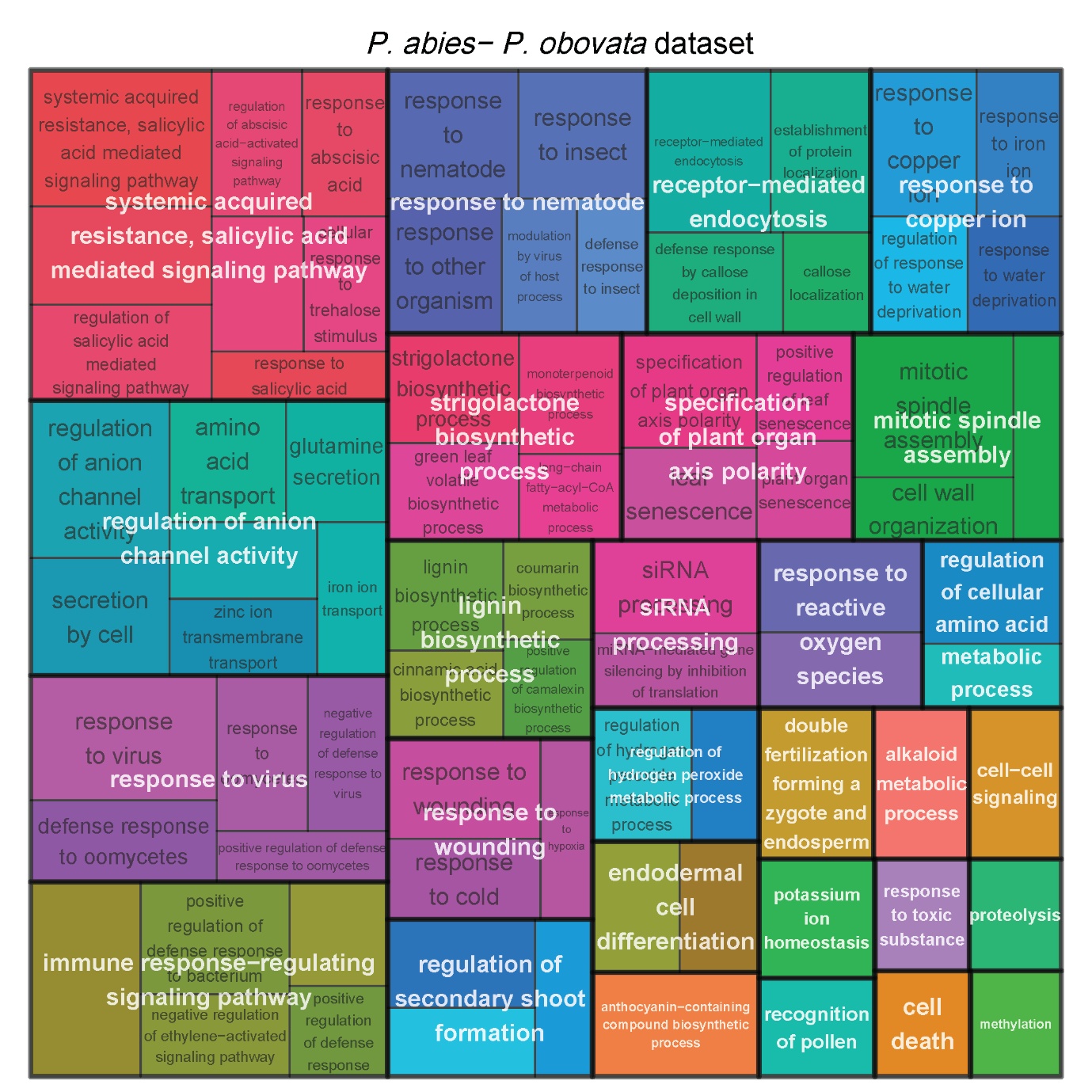

**Figure S3.** Reduced GO term (biological process) enrichment for the gCNVs identified in the *P. abies-P. obovata* dataset (*p*-value < 0.05).

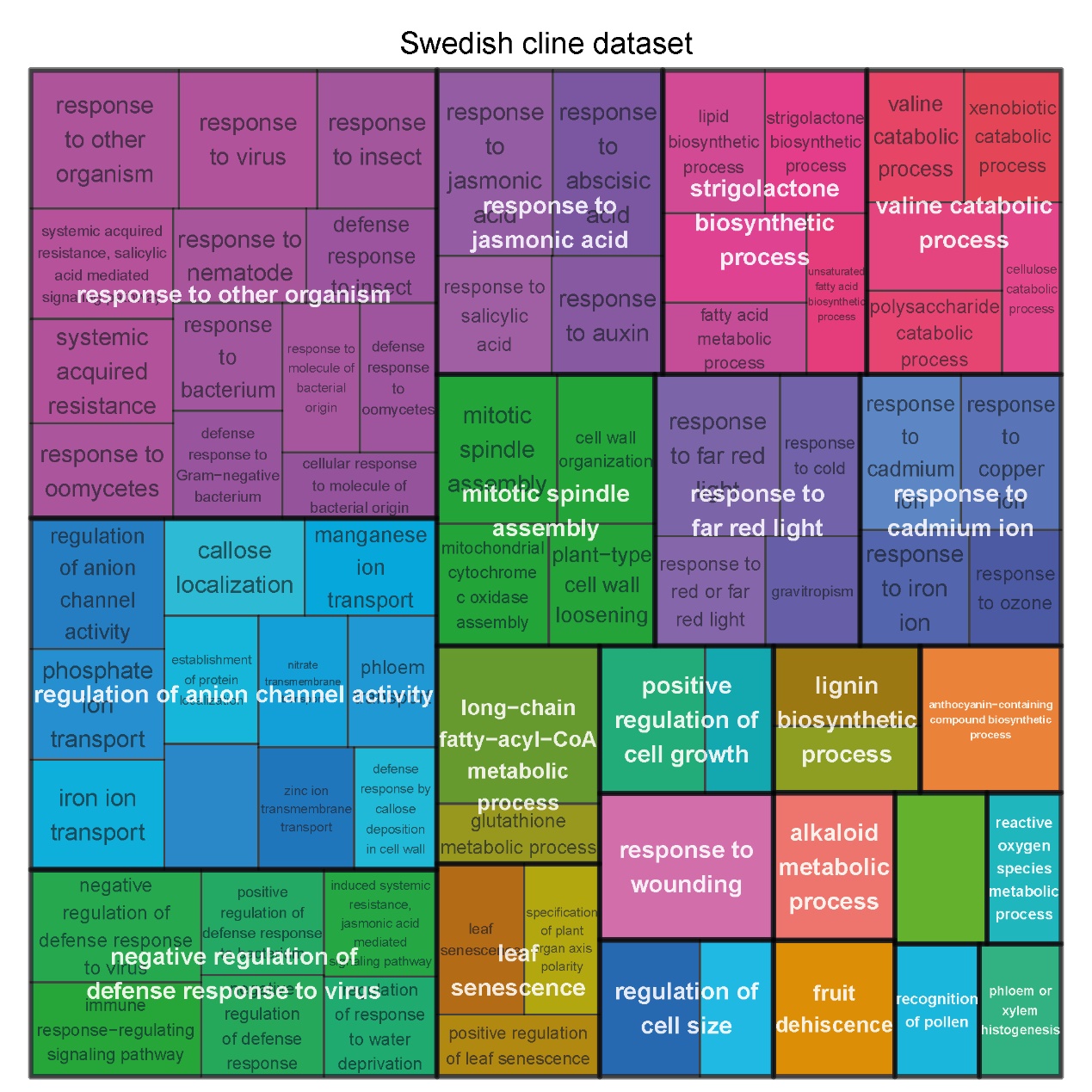

**Figure S4.** Reduced GO term (biological process) enrichment for the gCNVs identified in the *Swedish cline* dataset (*p*-value < 0.05).

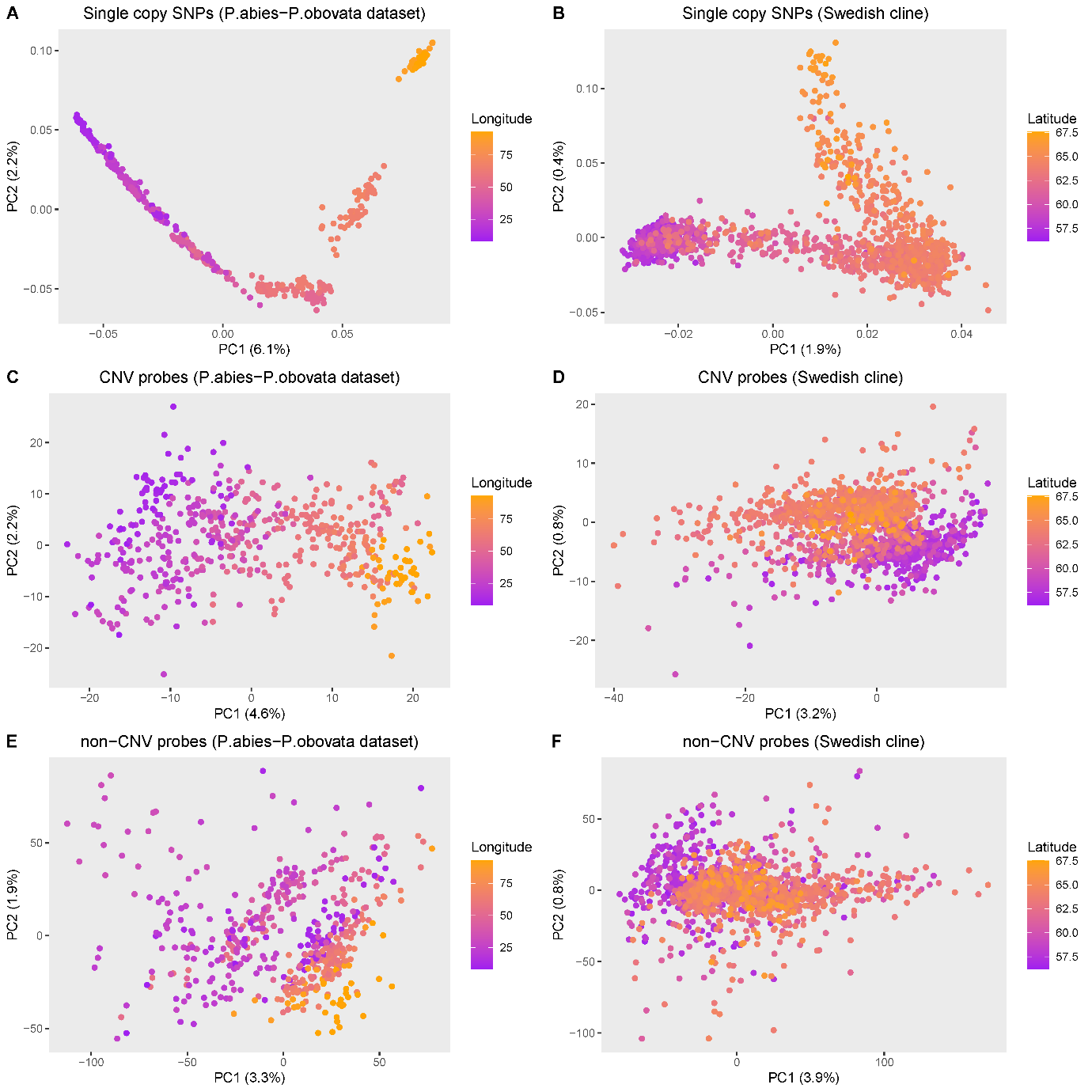

**Figure S5.** PCA analyses using either SNPs (top row), normalized depth of coverage of gCNVs (middle row) or normalized depth of coverage of single-copy genes (bottom row), for the *P. abies-P .obovata* dataset (left column) or the *Swedish cline* dataset (right column). Color gradients correspond to geographic gradient.

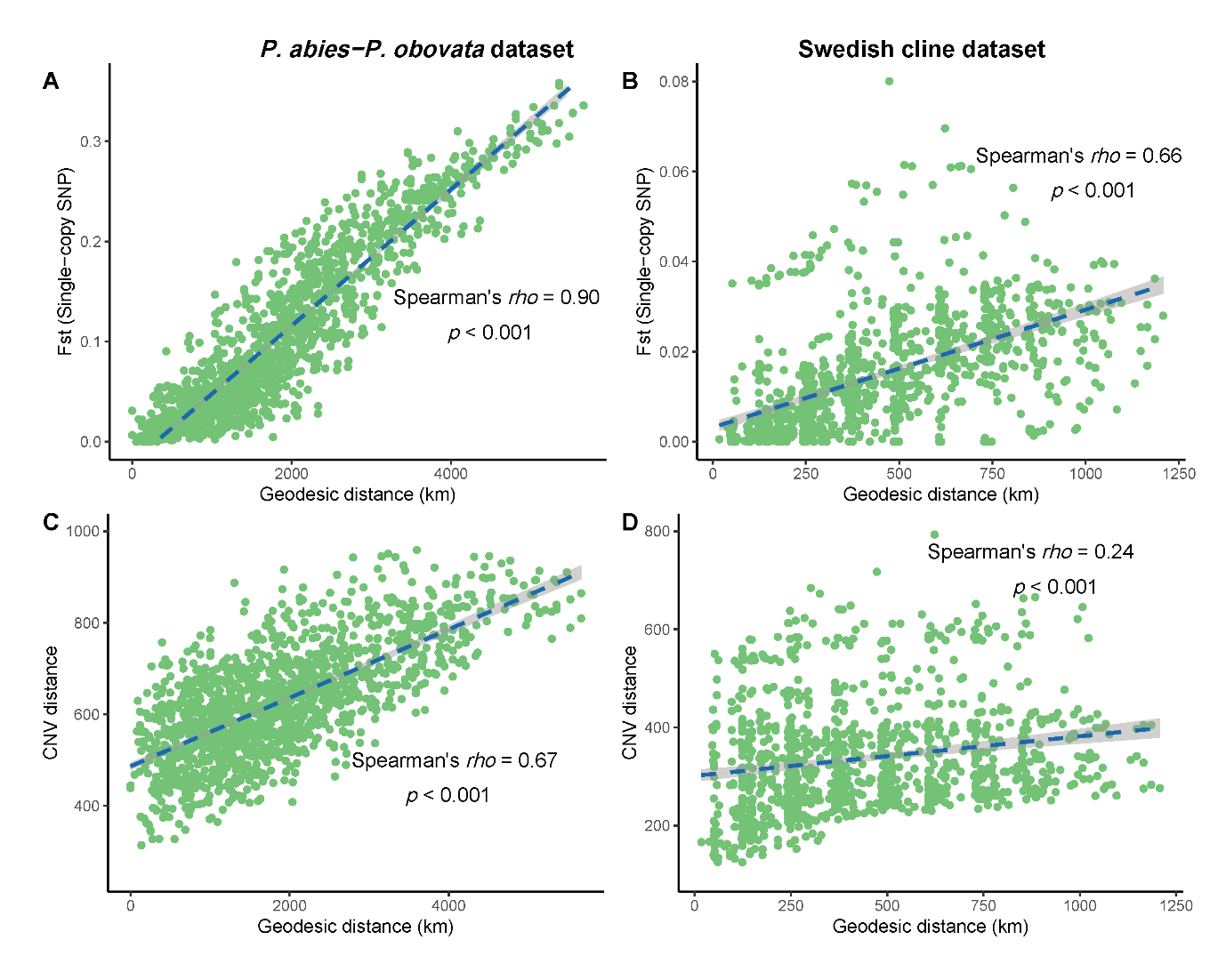

**Figure S6.** Patterns of Isolation by Distance using either SNPs (top row) or gCNVs (bottom row) for the *P. abies-P .obovata* dataset (left column) or the *Swedish cline* dataset (right column).

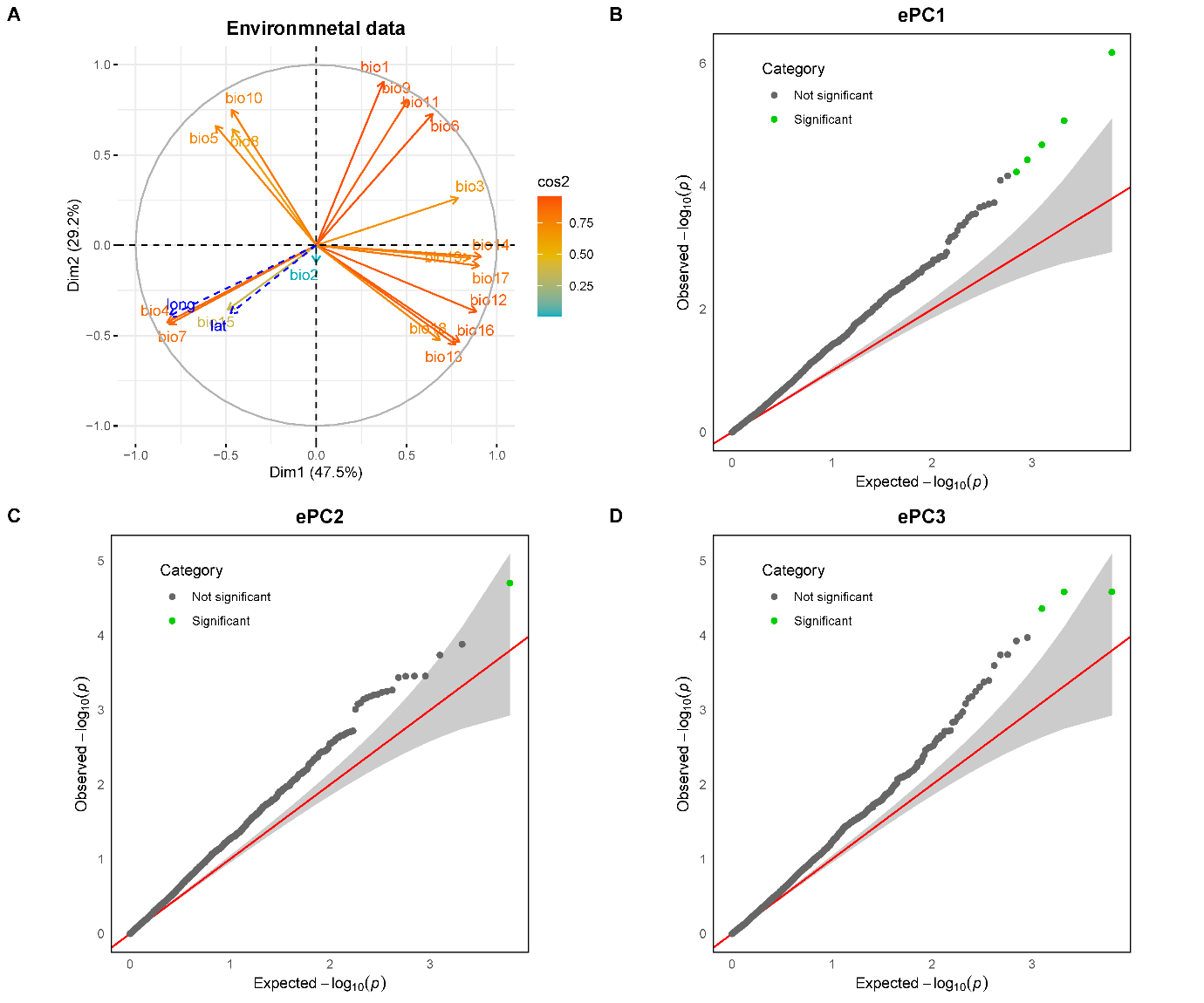

**Figure S7.** Genotype–environment associations in the *P. abies-P. obovate* dataset. **A**, The relative contribution of each of the 19 bioclimatic variables used in the PCA is indicated by the length and color intensity of the arrows. **B**, **C & D**, Q–Q plots for GEA between gCNVs and each of the three top PCs of the bioclimatic variables-based PCA. Grey shading indicates the 95% confidence interval under the null expectation (red line), and significant associations (adjusted *p*-value < 0.2) are in green.

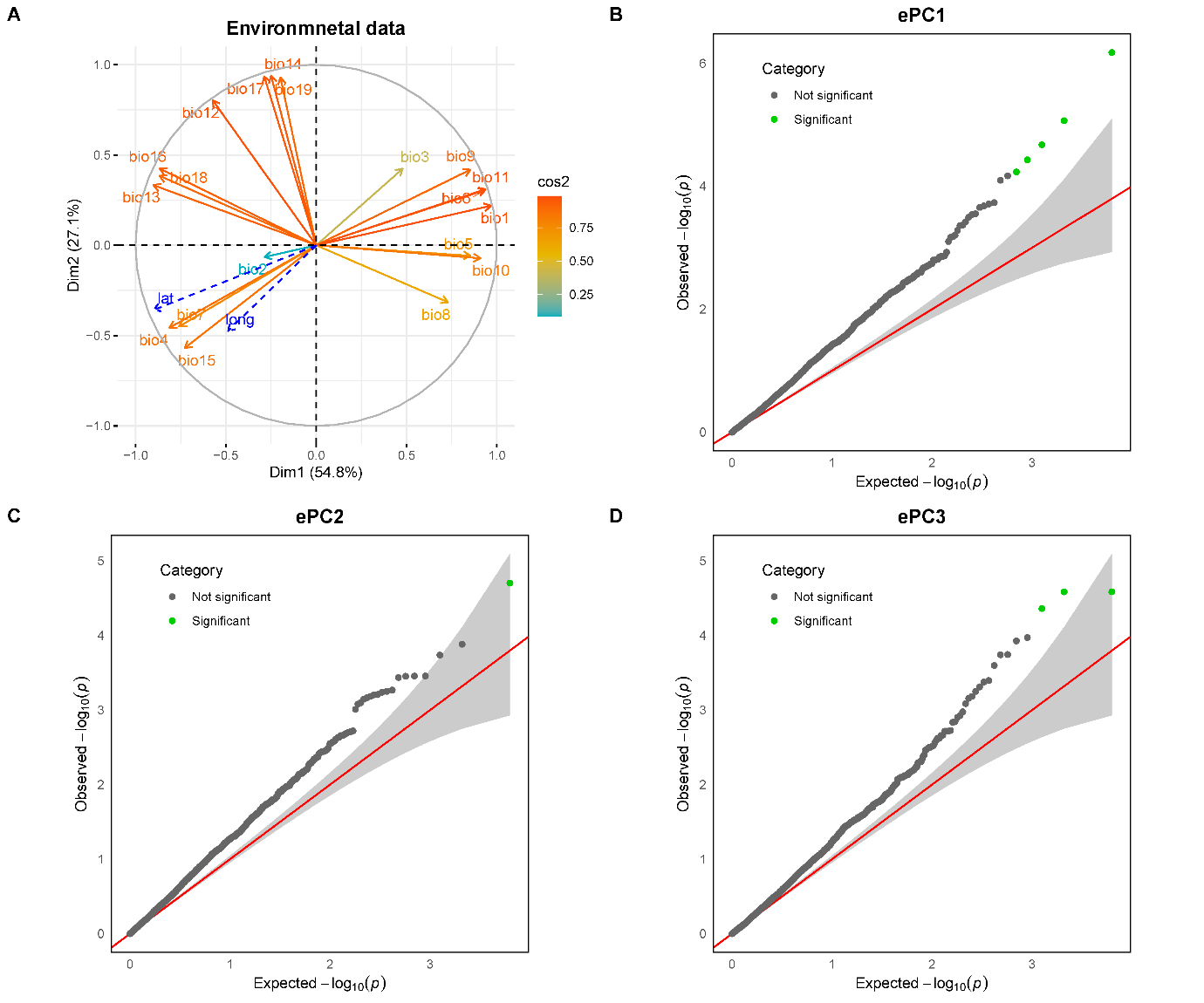

**Figure S8.** Genotype–environment associations in the Swedish cline dataset. **A**, The relative contribution of each of the 19 bioclimatic variables used in the PCA is indicated by the length and color intensity of the arrows. **B**, **C & D**, Q–Q plots for GEA between gCNVs and each of the three top PCs of the bioclimatic variables-based PCA. Grey shading indicates the 95% confidence interval under the null expectation (red line), and significant associations (adjusted *p*-value < 0.2) are in green.

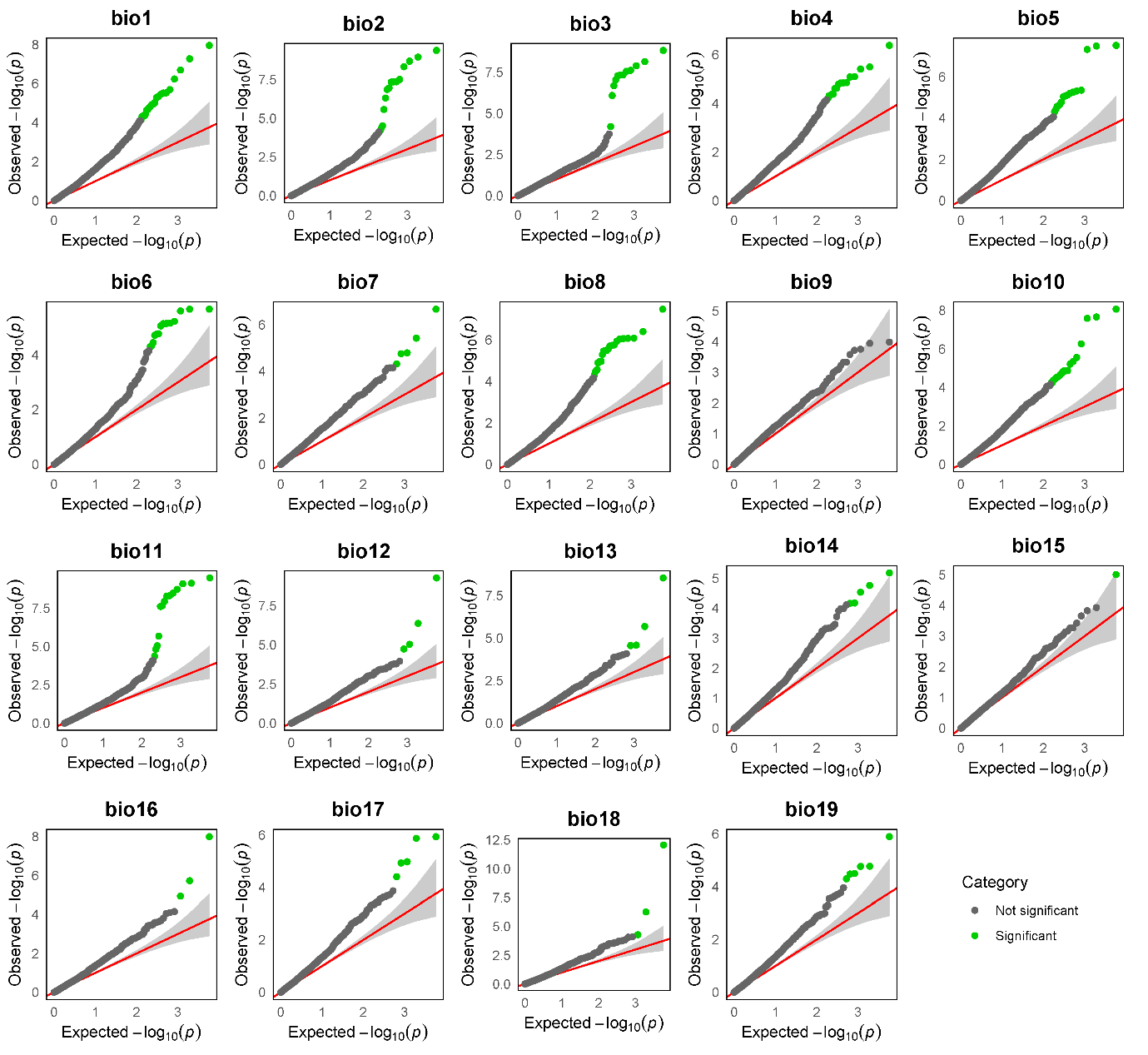

**Figure S9.** Q–Q plots of observed versus expected -log10 *p*‐values for GEA between normalized DoC of gCNVs and the each of the 19 bioclimatic variables for the *P. abies-P. obovata* dataset. Grey shading indicates the 95% confidence interval under the null expectation (red line), and significant associations (adjusted *p*-value < 0.2) are highlighted in green.

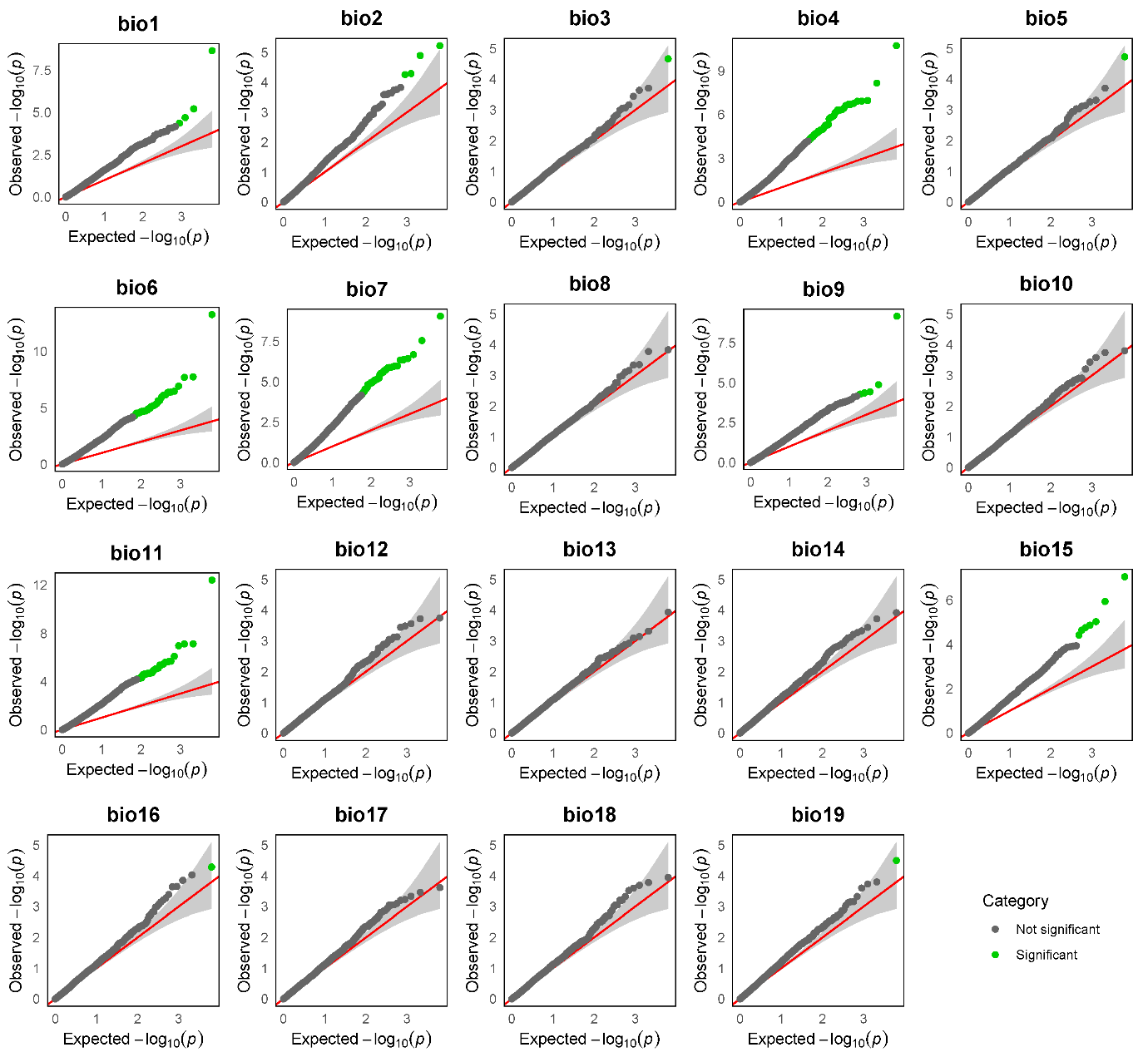

**Figure S10.** Q–Q plots of observed versus expected -log10 *p*‐values for GEA between normalized DoC of gCNVs and the each of the 19 bioclimatic variables for the *Swedish cline* dataset. Grey shading indicates the 95% confidence interval under the null expectation (red line), and significant associations (adjusted *p*-value < 0.2) are highlighted in green.

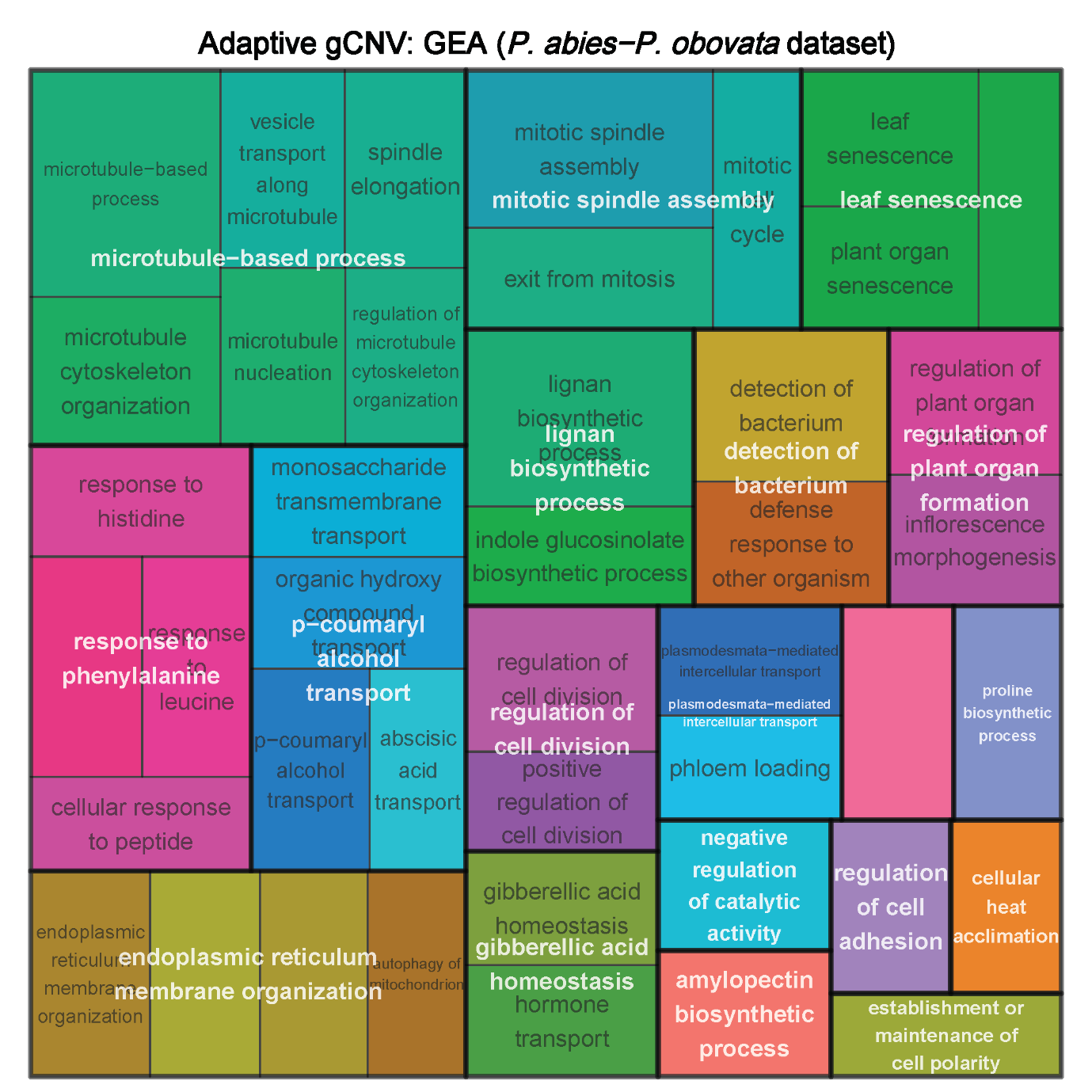
**Figure S11.** Reduced GO term (biological process) of significant enrichment (*p*-value < 0.05) for putatively adaptive gCNVs in the *P. abies-P. obovata* dataset identified using GEA.

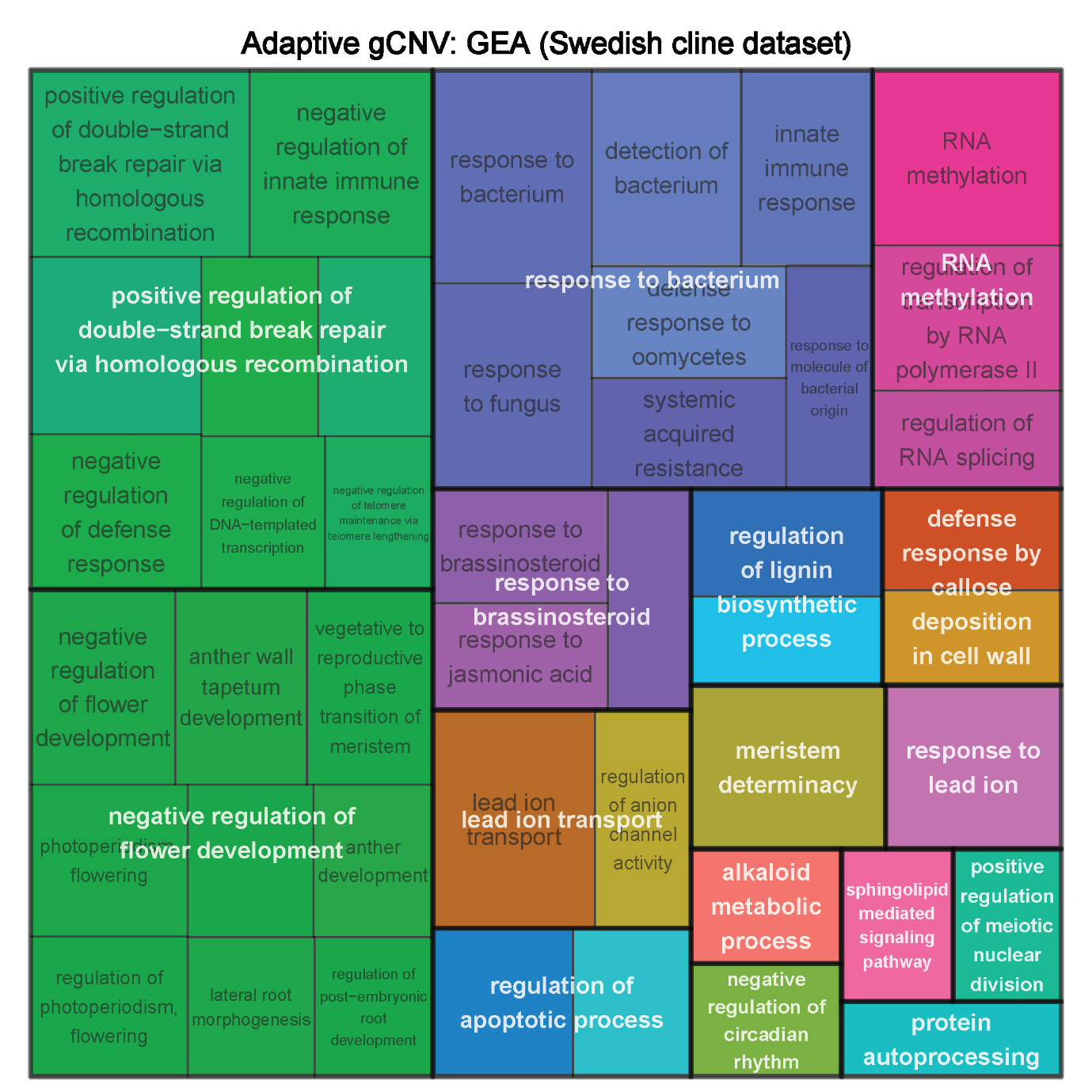

**Figure S12.** Reduced GO term (biological process) of significant enrichment (*p*-value < 0.05) for putatively adaptive gCNVs in the *Swedish cline* dataset identified using GEA.

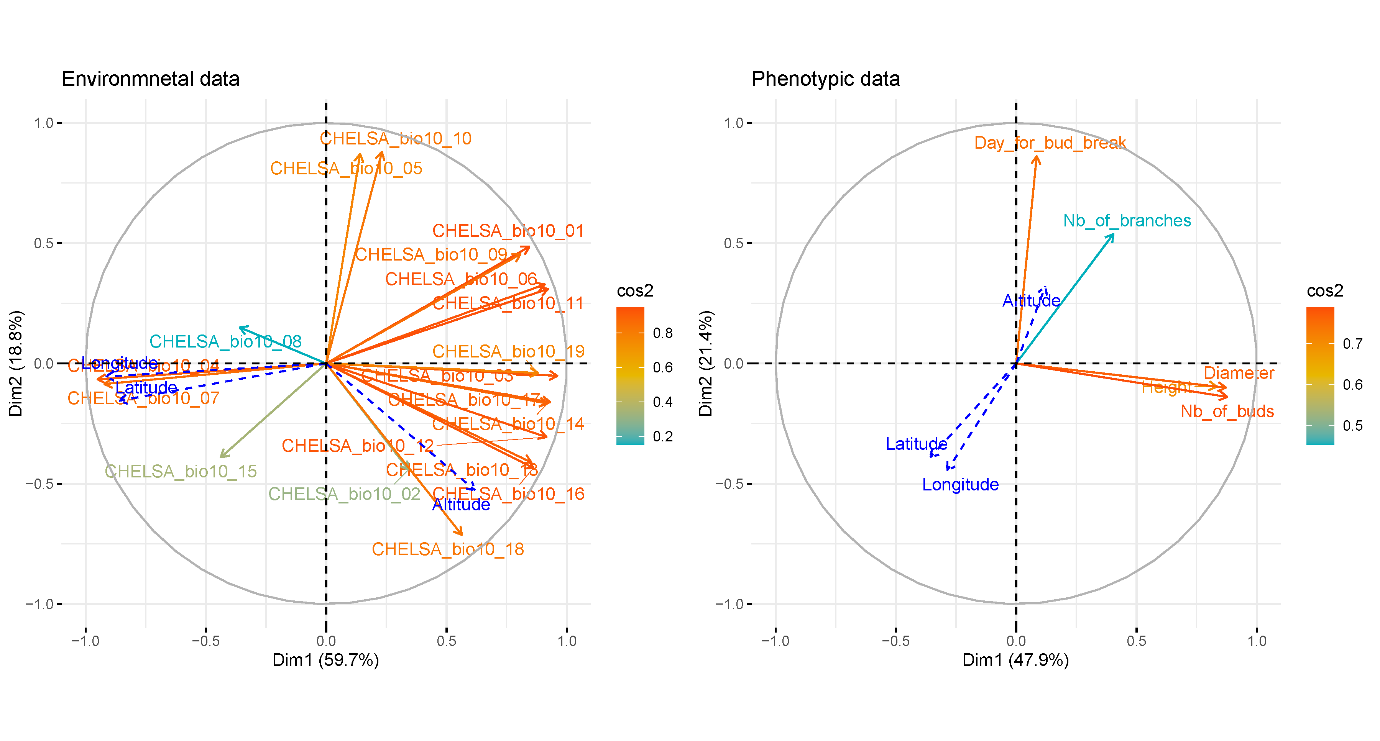

**Figure S13.** Principal component analyses (PCAs) based on 19 bioclimatic variables at tree sampling location, left, or phenotypic traits, right, for 230 individuals grown in controlled conditions; blue dashed arrows are posterior projections of latitude, longitude and altitude at sampling locations. The color gradient represents the relative contribution of the variables to the PCs.

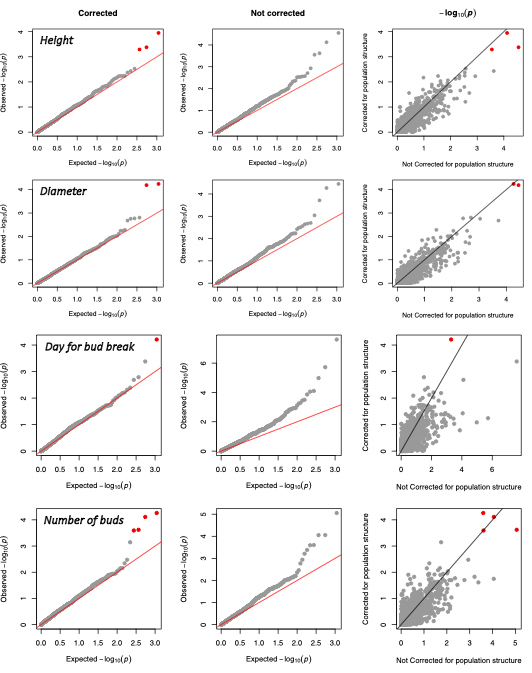

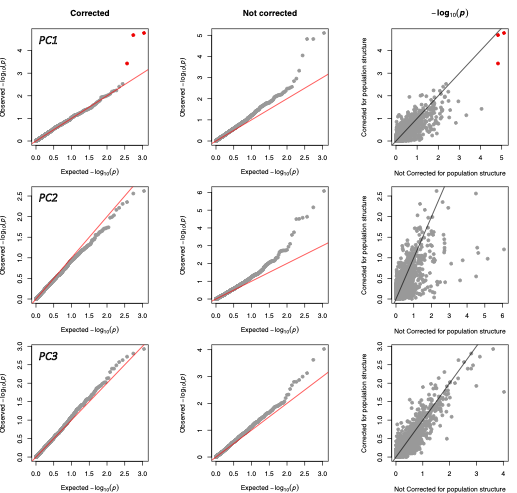

**Figure S14.** Gene copy number based GWAS. Each row corresponds to a phenotypic trait or a PC from a PCAs based on the phenotypic traits. QQ-plots present the observed -log_10_(*p*-value) as function of the expected ones for a GWAS controlling (left panels) or not (middle panels) for the population structure. The right panels give the correlation between the observed *p*-values controlling (y-axis) or not (x-axis) for population structure, the solid black line is the one-to-one line. An association was considered significant if satisfying the following criteria: FDR corrected *p*-value < 0.2 when controlling for population structure and < 0.1 when not controlling for it. Significant associations are indicated in red.

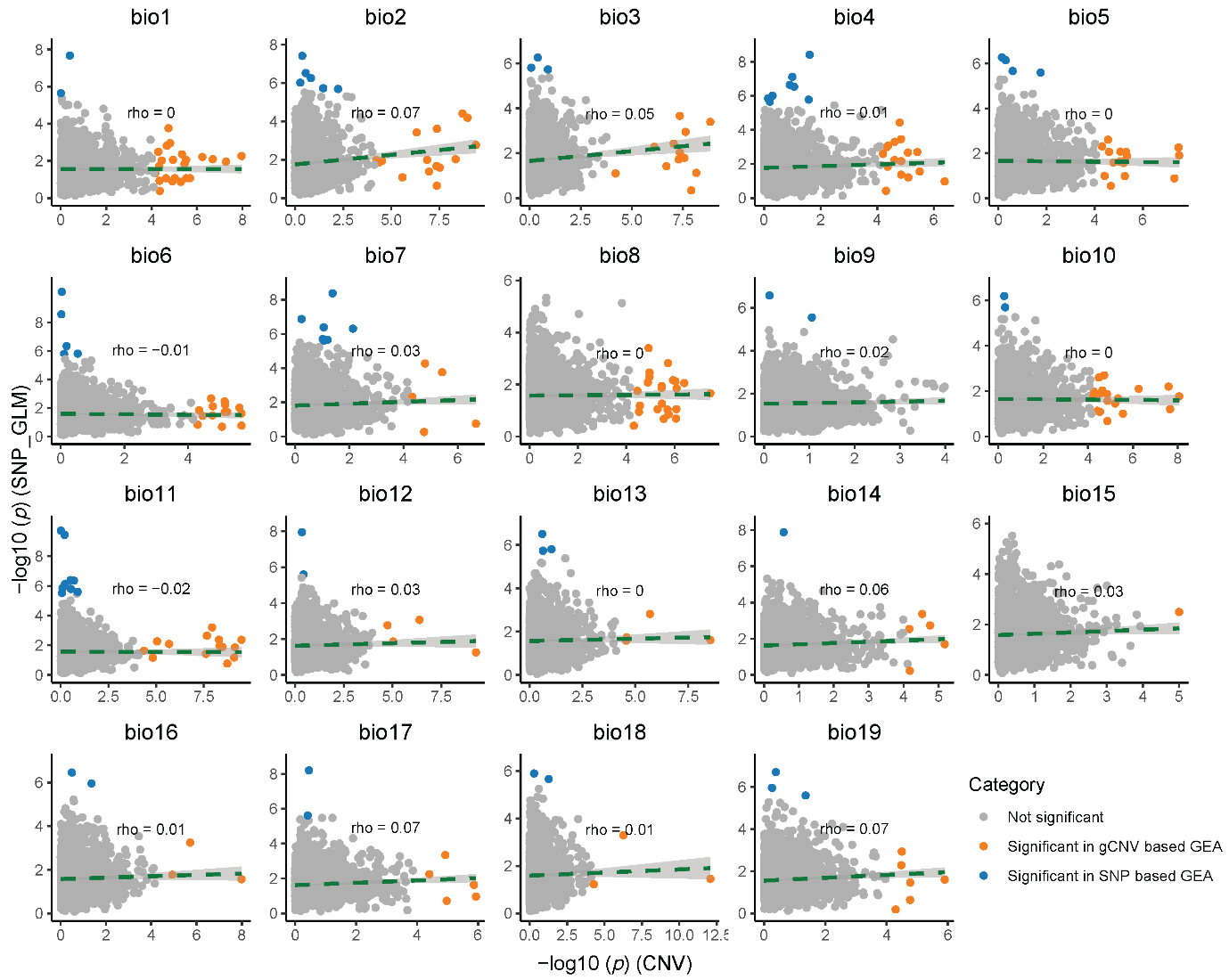

**Figure S15.** Relationships between gCNV- and SNP-based association across environmental variables for the *P. abies-P. obovata* dataset. x-axis: –log₁₀ *p*-values from gCNV-based GLM; y-axis; –log₁₀ of the smallest *p*-value of the SNPs within each CNV-probe. Significant probes in gCNV-based GEA and probes containing at least one significant SNP in SNP-based GEA are indicated in orange and blue, respectively.

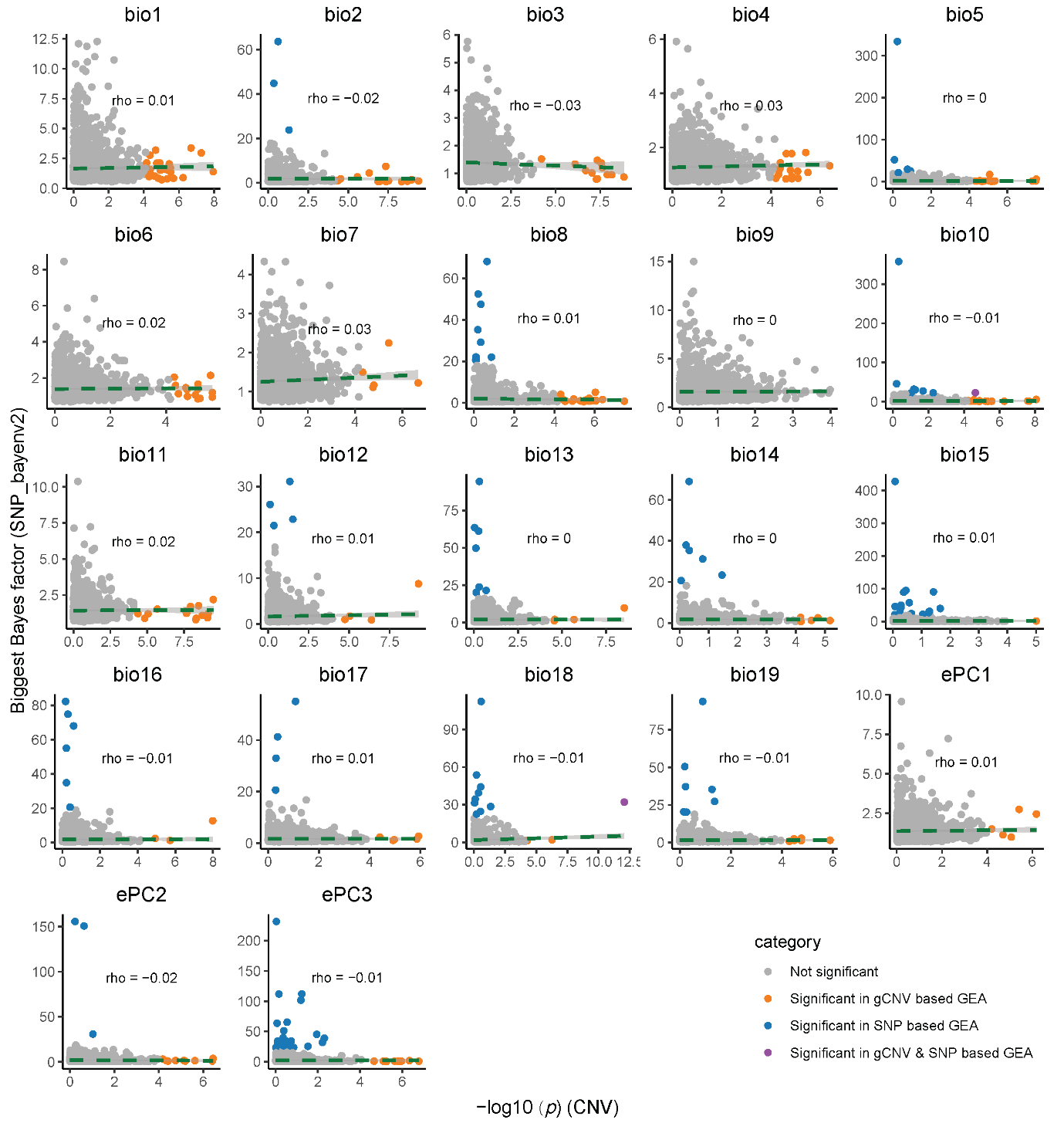

**Figure S16.** Relationships between gCNV- and SNP-based association signals across environmental variables for the *P. abies-P. obovata* dataset. x-axis: –log₁₀ *p*-values from gCNV-based GLM; y-axis; hisghest Bayes factor obtained with *bayenv2* of the SNPs within each CNV-probe. Significant probes in gCNV-based GEA and probes containing at least one significant SNP in SNP-based GEA are indicated in orange and blue, respectively, and in violet if significant in both.

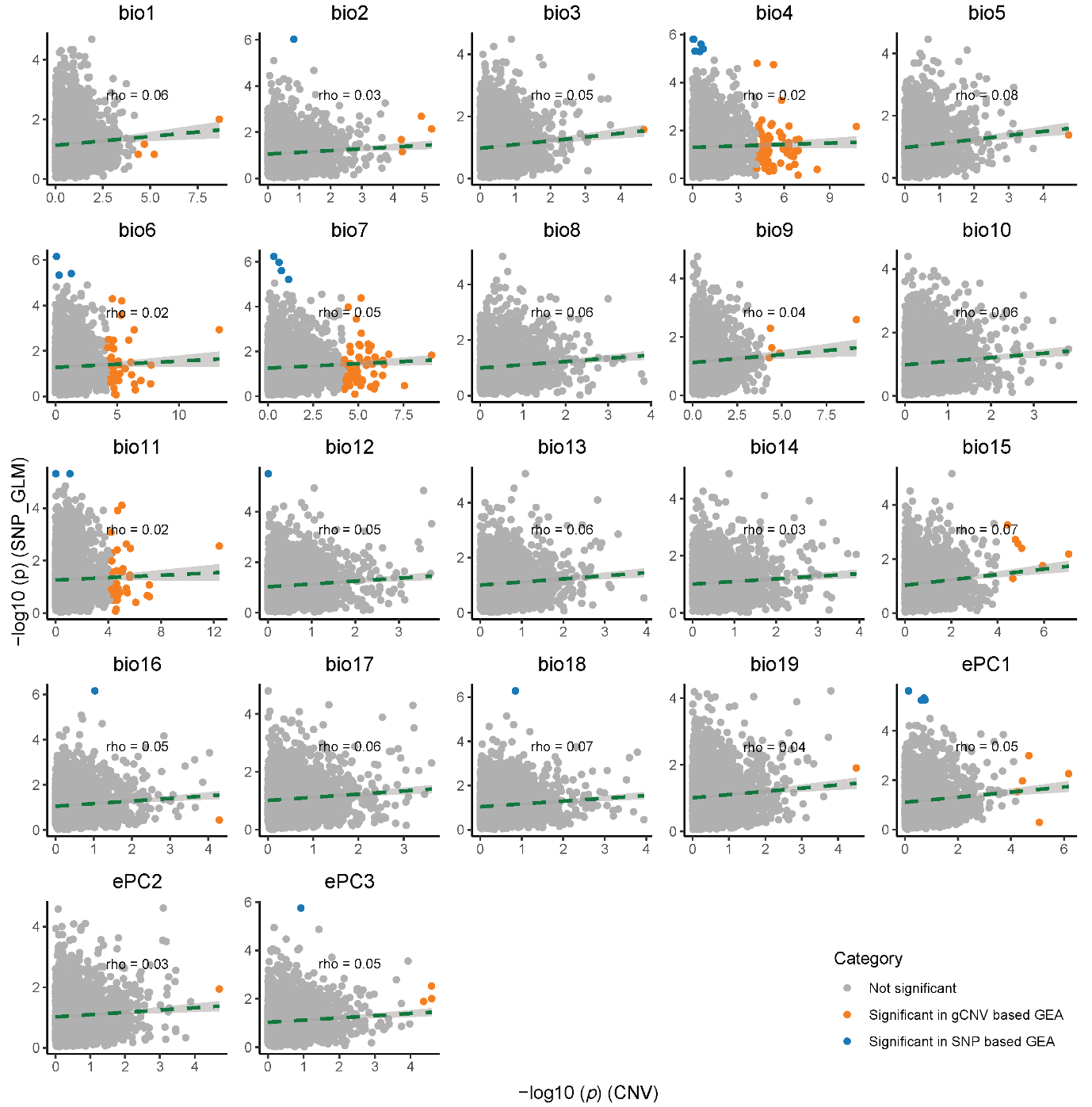

**Figure S17.** Relationships between gCNV- and SNP-based association across environmental variables for the *Swedish cline* dataset. x-axis: –log₁₀ *p*-values from gCNV-based GLM; y-axis; –log₁₀ of the smallest *p*-value of the SNPs within each CNV-probe. Significant probes in gCNV-based GEA and probes containing at least one significant SNP in SNP-based GEA are indicated in orange and blue, respectively.

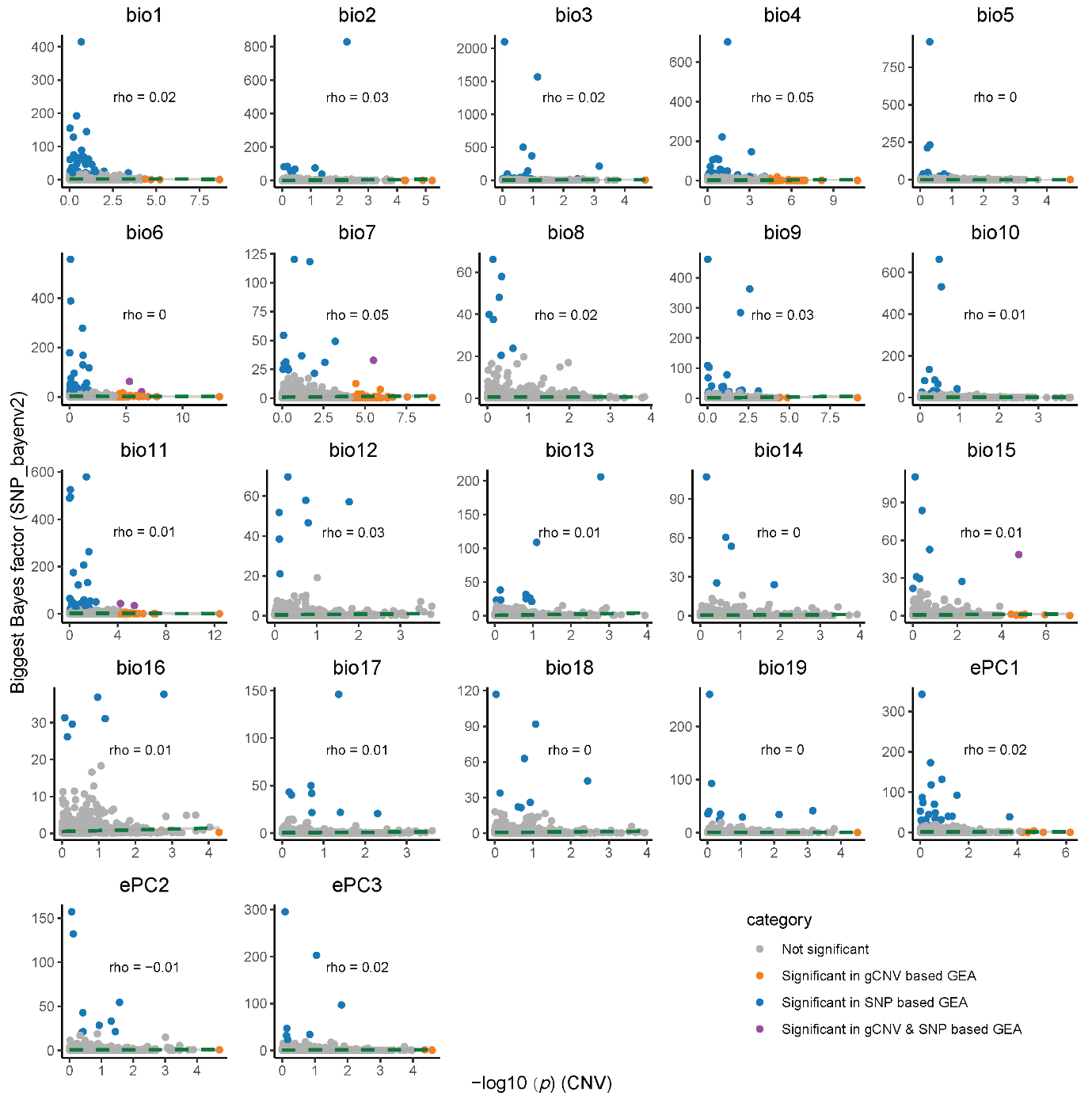

**Figure S18.** Relationships between gCNV- and SNP-based association signals across environmental variables for the *Swedish cline* dataset. x-axis: –log₁₀ *p*-values from gCNV-based GLM; y-axis; highest Bayes factor obtained with *bayenv2* of the SNPs within each CNV-probe. Significant probes in gCNV-based GEA and probes containing at least one significant SNP in SNP-based GEA are indicated in orange and blue, respectively, and in violet if significant in both.

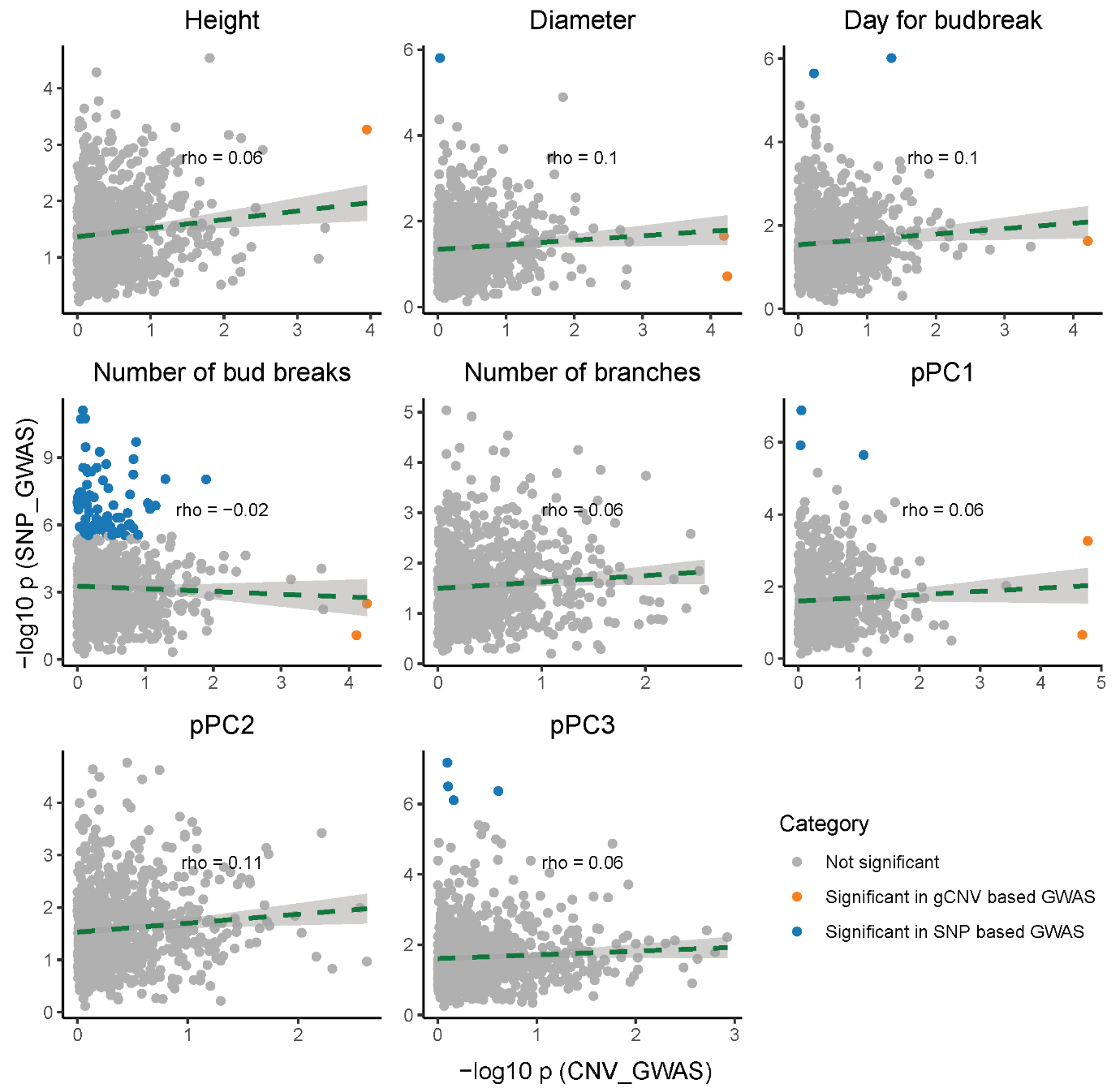

**Figure S19.** Relationships between gCNV- and SNP-based GWAS signals between five traits and top three principal components (pPCs). x-axis: –log₁₀ *p*-values from gCNV-based GWAS; y-axis; –log₁₀ of the smallest *p*-value of the SNPs within each probe from SNP-based GWAS.
